# Neural representation of association strength and prediction error during novel symbol-speech sounds learning

**DOI:** 10.1101/2023.11.06.564575

**Authors:** Gorka Fraga-González, Patrick Haller, David Willinger, Vanessa Gehrig, Nada Frei, Silvia Brem

## Abstract

Efficient learning of letters-speech sound associations leads to specialization of visual and audiovisual brain regions and is necessary to develop adequate reading skills. We still do not understand the brain dynamics of this learning process, and the involvement of learning and performance monitoring networks is still underexplored. Here we examined a feedback learning task with two mutually dependent parts in which novel symbol-speech sound associations were learned by 39 healthy adults. We used functional magnetic resonance (fMRI) and a reinforcement learning drift diffusion model that described learning across trials. The model-based analysis showed that posterior-occipital activations during stimulus processing were positively modulated by the trial-by-trial learning, described by the increase in association strength of each audiovisual pair. Prediction errors, describing the update mechanism to learn with feedback across trials, modulated activations in several mid-frontal, striatal and cingulate regions. The two task parts yielded a similar pattern of results although they varied in their relative difficulty. This study demonstrates which processes during audiovisual learning contribute to the rapid visual specialization within an experimental session and delineates a set of coactivated regions engaged in learning from feedback. Our paradigm provides a framework to advance our understanding of the neurobiology of learning and reading development.

## 1 Introduction

Learning to map symbols to speech sounds is an important milestone in learning to read alphabetic orthographies. This process leads to reorganization of visual and multisensory integration areas when learning how to read ^1,2^. Brain activity in these areas correlate with reading skills and developmental dyslexia has been associated with atypical development and, mainly, reduced activation in these regions compared to unimpaired readers ^3–5^. A crucial aspect which is still unclear is what other learning-related system contribute to the development of an efficient reading network during the acquisition of reading skills ^6^. In addition, there is evidence suggesting a potential involvement of frontostriatal circuits in reading disorders ^7–9^ (but see ^10^. To better understand such contributions will be an important step forward to characterize the neurobiology of reading and learning disorders.

Previous neuroimaging studies in this context have examined how children develop well-established letter-speech sound associations over months or years of instruction ^11–13^, or after trainings or interventions over several days or weeks ^14–16^. A previous study on pre-reading children following a short training (< 40 min) of artificial letters suggested that temporo-parietal responses to the trained symbols could be used to predict future reading skills ^17^. The evidence from these studies demonstrate that visual and multisensory areas specialize after varying learning period durations and in different developmental stages. Importantly, several behavioral studies support the relation of this type of novel symbol-sound learning and different early reading and cognitive skills in kindergarteners ^18^, 5-year olds ^19^ and adults ^20^. However, they still lack specificity in terms of cognitive mechanisms and additional brain systems supporting this specialization of visual and audiovisual areas. In addition, one of the studies suggests a narrow time window for detecting specialized responses^11^, which calls for a closer examination of learning dynamics within a smaller time frame.

In order to avoid over-learned and over-exposed stimuli, many studies in adults used artificial script learning tasks, and showed increased ventral occipito-temporal activations after few days of training ^21–23^. A recent study using magnetoencephalography (MEG) examined brain activations in adults when learning novel grapheme-phoneme associations and monitored performance during the learning block as well as one day after learning ^24^. The study included training blocks in which the audiovisual pairs were followed by learning cues, and test blocks in which participants had to respond whether a given pair was a match, no match or unknown. Besides learnable pairs, there were control pairs for which the learning cues were not informative. The results showed changes in superior temporal and dorsal parietal sources with learning. In addition, middle and inferior temporal regions, possibly reflecting activation from regions like insula or hippocampus, were engaged when using the cues to learn associations. This MEG study, although limited in its spatial resolution, provides an interesting window into rapid dynamics within early grapheme-phoneme learning.

Two previous EEG studies used a similar learning task to examine differences between typical and dyslexic adult readers in physiological responses to feedback ^25^ and in oscillatory networks ^26^. Their findings suggested differences between typical and impaired readers that would require further specification both in terms of brain networks and cognitive processes involved. In the current study we used a feedback learning task (FBL) adapted from those studies. The task required participants to learn associations between unfamiliar false fonts (i.e., letter-like symbols) and speech sounds based on the feedback presented on screen after a yes/no response. In this paradigm, learning occurs within the same block where participants respond, and brain activations related to learning novel audiovisual pairs are examined on a trial-by-trial basis within three experimental blocks (< 8 min each). Our task was set to simulate an important part of learning an alphabetic script, where practice and trial-errors and feedback allow the reader to establish new associations. In addition, we included an additional task part that simulates the role of diacritic marks, common in orthographic languages, as well as the inconsistencies between phonemes and graphemes, as in opaque orthographies like English. This additional task part (FBL-B) follows the same principles and depends upon the stimuli from the main part (FBL-A), since diacritic marks modify the speech sounds associated with the false fonts from the preceding FBL-A blocks. The additional goal of part B, which is expected to be more demanding, was as well to generate an additional level of difficulty and variability in performance. The adult population in this study was also chosen to allow flexibility in length and complexity of the paradigm design, which is important for our goal of finding new neural and behavioral descriptors of individual variability in learning.

The previous work from Fraga Gonzalez et al., (2019, 2022) lacked specificity in describing the brain areas involved in this task. The current functional magnetic resonance imaging (fMRI) environment allows a spatially-resolved network characterization not possible with electrophysiological recordings. Here, the frontostriatal circuits are of special interest. Activity in the anterior cingulate cortex (ACC) has been associated with a variety of functions relevant to our task, like feedback/reinforcement learning, error detection, action selection and conflict monitoring (see review in ^27^). A recent review argues for three core computational principles in the ACC: hierarchical decision making, spatiotemporal models of the environment and cost evaluation ^28^. In this context, an important concept is the prediction error (PE), that is the discrepancy between expectations and outcomes, proposed to drive learning ^29^. The ACC is involved in adjusting responses and decision making using this predictive signal together with the surrounding prefrontal cortex ^30^. The regions involved in encoding these PE signals are striatal regions like caudate and putamen ^31,32^, which are also linked to different forms of motor, instrumental and associative learning ^33,34^. In the context of reading acquisition, this predictive coding framework has been used to explain the trajectories in specialization of visual occipito-temporal regions for orthography ^35^.

In order to describe these learning components, our experimental task was analyzed with a reinforcement learning drift diffusion (RLDD) model that allows describing both trial-by-trial learning as well as the global dynamics of decision making in terms of speed/accuracy trade-offs ^36^. The choice of this model broadens the cognitive descriptors that can be derived, and makes it an interesting tool to search for novel markers that could ultimately help characterizing clinical populations. The model-based cognitive neuroscience approach in this analysis is intended to capture underlying cognitive processes and their associated brain activation patterns which may be overlooked in a basic reaction times and accuracy analysis ^37^.

In summary, the primary goal of the current study is to characterize changes in brain with trial-by-trial learning of symbol-speech sound associations, with a focus on visual/audiovisual regions and regions involved in associative learning and feedback processing. As a secondary aim, the current paradigm targets individual differences in learning that could be linked to cognitive performance and reading skills. Ultimately, the broader goal of this study is to search for new neurocognitive markers to predict and characterize typical and (a)typical reading development.

## 2 Methods

### 2.1 Participants

The current study is based on a sample 39 healthy adult participants (21 females; age 25.19 ± 3.12 [range 18.14-32.51], see Table 1 for details. Participants were recruited using University platforms and social media to be right-handed (Swiss-) German-speaking and between the age of 18-35 years. They were screened for contraindications for MRI (e.g., metallic implants, neurostimulators or cardiac pacemakers, pregnancy) and neurological disorders. Two of participants in the sample reported attentional problems and problems with spelling and reading but no diagnosis of psychiatric disorders. The current study sample was obtained from an initial pool of 43 participants from which 1 participant was excluded due to voluntary interruption of the scanning, 1 was excluded due to technical problems during scanning, 1 participant did not comply to the experimental task instructions and 1 participant could not be scanned due to MR compatibility issues. A further exclusion criteria was a non-verbal IQ < 80 in the cognitive assessments, not fulfilled by any participant in this analysis. Additionally, participants filled in an adult reading history questionnaire ^38^ on their reading history and habits. The project was approved by the local ethics committee of the Canton of Zurich in Switzerland and was performed in accordance with the declaration of Helsinki. Participants signed a written informed consent form before participating in the study.

**Table 1.**
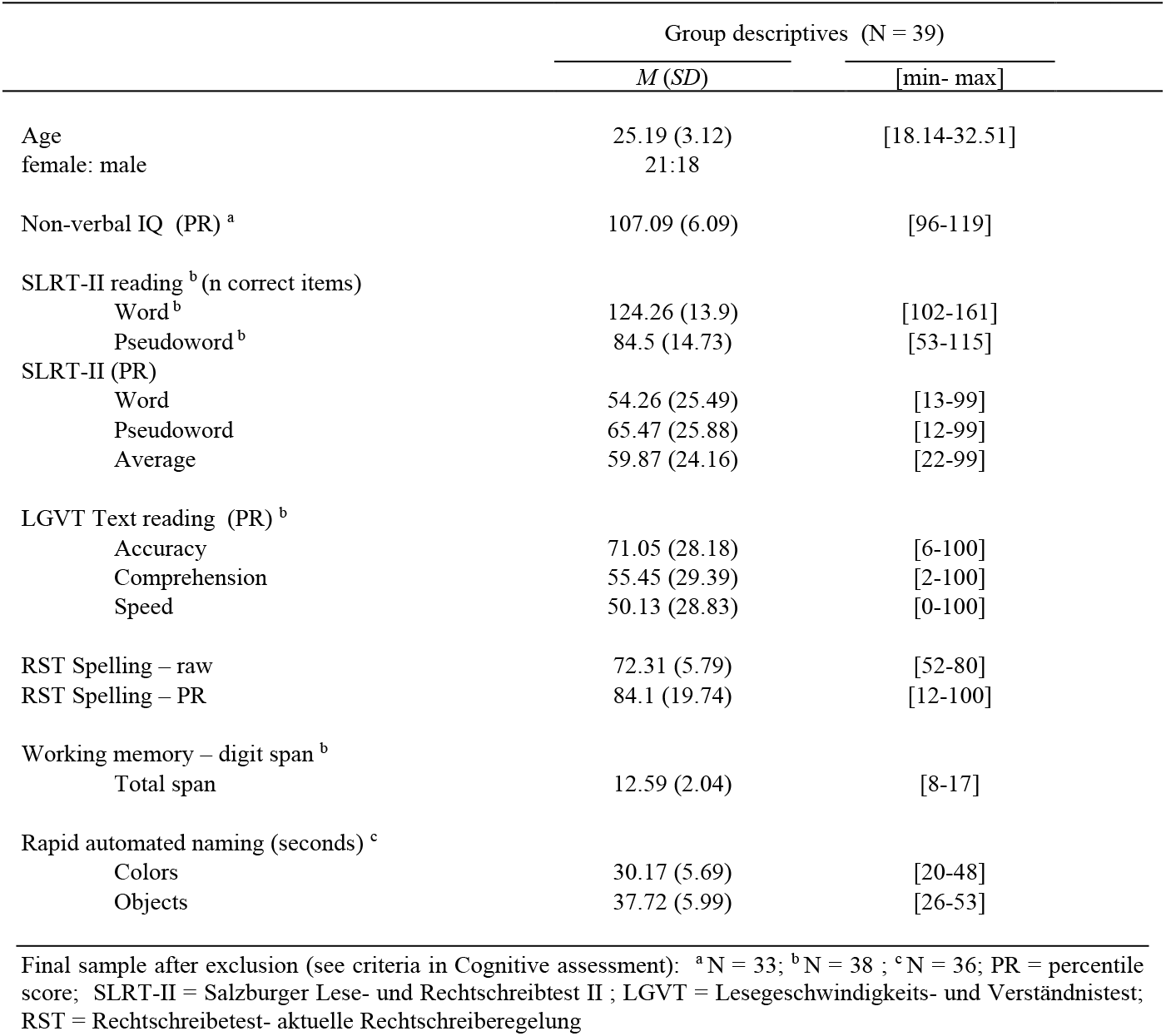
Descriptive statistics showing cognitive performance.

### 2.2 Cognitive assessments

All participants performed a series of cognitive and reading tests. The descriptive statistics of performance are summarized in Table 1.

Due to Covid-19 related measures some of the tests were conducted via video-call using an application secured by the university (https://uzh.meet.switch.ch/). The following tests were conducted online. Working memory was assessed with the backward and forward digit span subtest from the *Wechsler Adult Intelligence Scale-fourth edition (WAIS-IV;* ^39^). Rapid automated naming (RAN) was used as a measure of general naming speed with the object (animal) and color naming tasks from the *Test zur Erfassung der phonologischen Bewusstheit und der Benennungsgeschwindigkeit* (TEPHOBE; ^40^).

In addition, the following tests were performed in the MR facilities before commencing the neuroimaging recordings. Overt word and pseudoword reading fluency was assessed with the *Salzburger Lese- und Rechtschreibtest (*SLRT-II; ^41^). The number of correctly and overtly read items in one minute are used as main measure of this test. Since this is the main test to estimate reading abilities, the distributions of percentile scores are shown in Figure 1. Moreover, comprehension, velocity and accuracy of covert, text reading were assessed with the *Lesegeschwindigkeits- und Verständnistest für die Klassen 5-12+* (LGVT; ^42^). Participants are instructed to covertly read as fast and accurately as possible a brief text within 6 minutes. Within the text there are several single-choice questions in which they must select 1 out of 3 words that fit the paragraph content. The percentile scores are presented in Figure 1. Spelling was tested with the *Rechtschreibetest-aktuelle Rechtschreiberegelung* (RST; ^43^), in its short version for 14-60 years of age. In this test participants get a transcript of a text with some gaps that they must fill after listening to the experimenter read the text aloud. To avoid inconsistencies between experimenters we presented a video recording of a linguist reading the text. The non-verbal intelligence IQ was estimated with the *Reynolds intellectual assessment scales* (RIAS; ^44^).

**Figure 1.**
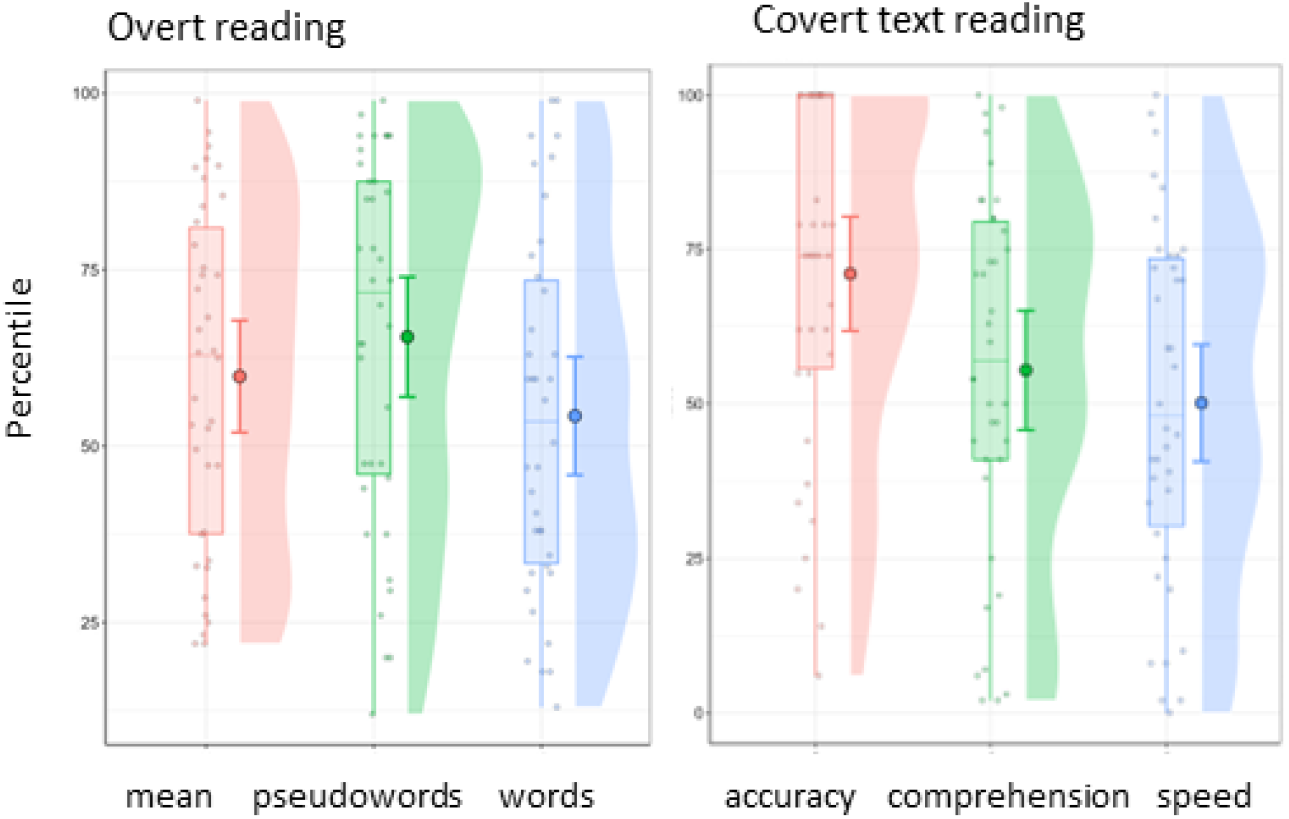
Percentile scores for overt reading scores (SLRT-II; left plot) and for covert, text reading accuracy, comprehension and speed scores (LGVT; right plot).

### 2.3 Task and stimuli

The task performed in the MR scanner is an adaptation and extension of a task previously used in two electrophysiological studies ^25,26^. It was programmed and presented using Presentation® software (version 20.1, www.neurobs.com). The task is illustrated in Supplementary Figure A.1. Participants were instructed they had to learn new symbol-speech sound mappings by deciding on each trial whether the audiovisually presented pair was correct or not, and then receiving feedback on the screen. In each trial, one symbol and one speech sound were presented simultaneously and participants were instructed to press left or right button to indicate whether they thought the pair was a correct match or no. They were instructed to respond in every trial and to guess if unsure in order to learn the associations via feedback. The left/right button assignment to yes/no responses was counterbalanced across participants. The duration of the trials was terminated by the participant’s button press and after each response, the feedback provided was a happy or a sad smiley to indicate if the response was correct or incorrect, respectively. If no response was given within a 2500 ms interval a ‘Schneller’ (‘Faster’) feedback would be presented before the next trial to prevent too slow responses or inactivity. All feedback remained for a mean of 2000 ms on screen (jittered durations drawn from a normal distribution of mean 2000 ± SD = 500 ms). A fixation cross followed feedback until the next trial. The durations of this fixation were drawn from a normal distribution of 2500 ± 500 ms. There were two learning blocks in each task, each block with 48 trials in which 6 different symbol-speech sounds could be learned. Each sound was repeated across 8 trials, 50 % of them showing the correct symbol and 50 % showing the incorrect symbol. Pseudorandomized sequences were generated constraining the appearance of consecutive trials in which the same pair or sound were presented.

The A and B tasks shared the same principles and design but varied in the stimuli presented (see Supplementary Figure A.1, panel B). They were presented always consecutively as the second part (B) builds on the stimuli learned in part A. In part FBL-A the false fonts were single characters from a pseudofont. In the subsequent FBL-B, three of the false fonts from the same block in part A were presented with two different diacritic marks on top, leading to six new false fonts associated with new speech sounds. The speech sounds were either completely different, a prolongation or a modification of those associated with the symbol in previous part. For example, the symbol N is paired with the speech sound /a/ in FBL-A and is presented as pairs P /aa/ and O /ao/ in FBL-B). The blocks in both parts started with a very brief practice sequences to ensure the participants understood the principle. In part A this practice showed 9 trials of 3 new false fonts and sounds that were not presented later in the block. In part B the practice included 6 response-terminated trials to ‘refresh’ the knowledge of the previously learned associations, by showing false fonts and sounds that would later be presented with the modifier marks.

### 2.4 Computational model

The basic analysis of task performance based on proportion of correct responses and reaction times (see statistical analysis) provide limited opportunities for interpretation on the underlying cognitive processes. Thus, we applied a computational model to derive more fine-grained parameters that can be later correlated with learning-related brain areas and with other cognitive skills. We chose a reinforcement-learning drift-diffusion model (RLDDM; ^36^) as it has been proposed to reveal both learning and decision making processes, which are both relevant to perform the task. There are several parameters from this model of special interest in our analyses. The parameter *association strength* from the reinforcement learning part of the model describes the process of mapping letters to speech sounds within the learning block. The main parameters of interest from the drift diffusion model, *drift rate* and *decision boundary*, describe how subjects accumulate information to make their responses, as well as trade-offs between speed and accuracy, respectively. In addition, the *non-decision time* parameter from the drift model disentangles the component from the reaction times that can be attributable to general processing speed rather than to a learning-based decision. The RLDDM is illustrated in Figure 2. The model parameters and priors are presented in Supplementary Table A.1.

**Figure 2.**
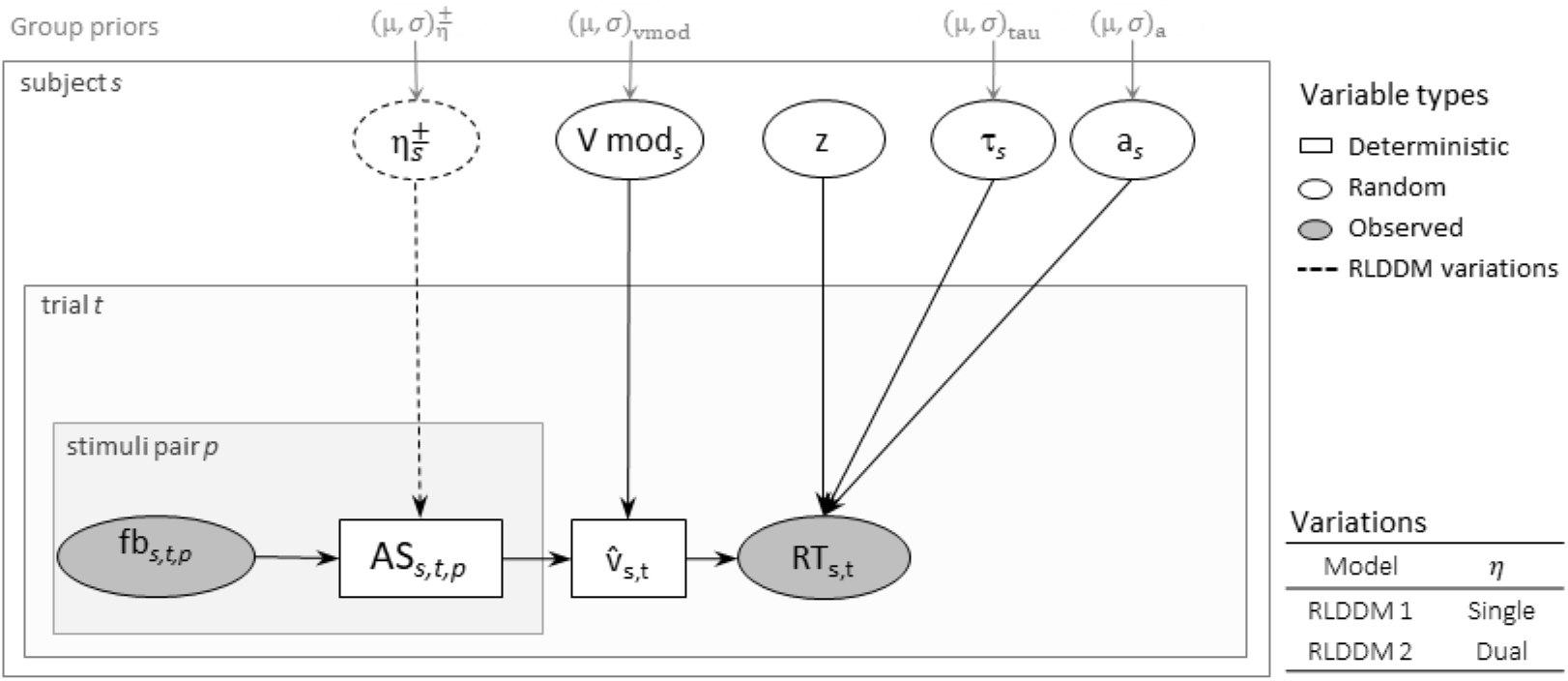
Schematic of the reinforcement learning drift diffusion model (RLDDM). Deterministic variables are illustrated with rectangular nodes, random variables with round nodes and observed data with grey round nodes. Parameters are estimated in a hierarchical Bayesian framework. The subject parameters are estimated from group means *μ* and variance *σ* parameters. Learning rates (*η*) modulate the trial-by-trial associative strength (AS; or expected value) which are updated depending on the feedback. The mean AS modulated by the *v* mod free parameter is used to estimate the drift rate *v* per trial. Learning rates could be single or dual (for + and – prediction errors). *a* = decision boundary; *τ* = non-decision time; *z* = starting-point (fixed parameter); fb = feedback; *s* = subject; t = trial; *p* = symbol-sound paired; v = drift rate

The first component of the RLDDM is a reinforcement learning model (RL; ^45^). The basic principle of RL models is that learning is driven by unexpected outcomes, like occurrence or absence of reward, which are captured by a prediction error, PE signal. The PE describes the difference between observed and predicted outcomes and is used to update expectations and adapt behavior in subsequent occurrences. This updating process is described by [1]

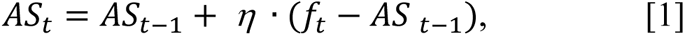

where *t* refers to the current trial. The formula describes how the association strength AS between the stimuli in a trial and its outcome are updated based on the differences between the feedback obtained *f* and the outcome expected from the previous occurrence. The term AS is used as equivalent to the expected values of an outcome ^46^. The trial-wise updating is modulated by a learning rate parameter *η* per subject, ranging from 0 to 1 and with larger values describing faster adaptation to grapheme-phoneme associations. In the RL models, the response choice is often modeled by using a softmax linking function ^47^ in which the probabilities on choosing a response over others are based on reward values. In the softmax case, the subject’s sensitivity to a reward is captured by a sensitivity parameter *β*, reflecting a subject’s individual importance of value differences. However, the softmax choice rule does not allow to take into account latency or differentiate between fast and accurate responses from those slow and conservative.

In the RLDDM, the softmax choice rule is replaced by a drift diffusion model (DDM; ^48^). The DDM model families assume that a decision between two options is based on accumulating noisy evidence in favor for one of the options, until a decision threshold is reached. These models have been applied to data from many different psychological domains (see an overview ^49^). In brief, in the DDM the distributions of accuracy and RTs depend on several parameters. The main parameter is the drift rate *v*, which indicates how fast the decision process reaches the boundaries (higher values mean faster and more accurate responses). The boundary separation or decision threshold parameter *a* adjusts how much noisy evidence is needed before reaching a threshold, i.e., the speed-accuracy trade off (higher values indicating slower and more accurate decisions). A starting point *z* would indicate an initial bias for a given response, this was set to 0.5 as the participant had no prior knowledge of the correct association. The non-decision time *τ* describes the time of encoding the stimuli and preparing the motor response. In the current RLDDM, the drift rate is described per trial *v*_*t*_, as the scaled mean of the AS of the two reinforced options. See formula [2]:

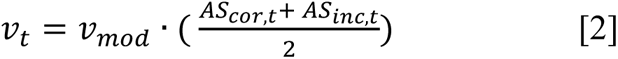

In our study, the expected values are defined as the AS of correct or incorrect responses. The scaling factor *v*_*mod*_ is a free parameter, similar to the inverse temperature *β* in the softmax rule, which describes the degree of sensitivity, that is, how much the choice is conditioned by association strengths. In the current analysis, the mean of the association strengths is used, since drift rate is expected to be highest when both associations are well-known, medium when only one is well-known and lowest when none of the associations has been learned. The RTs and accuracy are simultaneously estimated in RLDDM, as in the standard DDM, using the Wiener first passage time (WFPT) distribution. This distribution describes the likelihood of a observing a response and RT and uses the DDM parameters described above: [3]

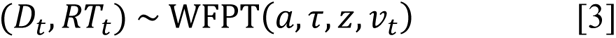

#### 2.4.1 Model fitting and model comparisons

Parameters were estimated using a Bayesian hierarchical modeling approach, which has been suggested to be advantageous for parameter estimation with limited amount of data ^50–52^. We used the Markov Chain Monte Carlo methods implemented in Stan (^53^, see also https://mc-stan.org/) with the interface to Stan, the ‘cmdstan’ package (version 2.25.0; ^54^). We ran 4 chains of 10000 iterations each with 4000 warm-up samples. Weakly informative priors were both modelled (see Supplementary Table A.1). Convergence was assessed with the Gelman-Rubin convergence diagnostic *R̂* ^55^. Values of *R̂* ≤1.01 are considered indicative of successful convergence; this criterion was fulfilled by our parameters (see trace plots of chains and convergence diagnostics in Supplementary Figure A.2.)

In the current study we run two model variations; one model with separate learning rates *ηs* for positive and negative PEs, and one with a single learning rate (^36^, see model illustration in Figure 2). We compared the models using the Widely Applicable Information Criterion (WAIC; ^56^). The WAIC is computed from the log pointwise predictive density using the variance of individual terms summed over the data points to correct for model complexity. The prior distributions from which the different group and subject level parameters were drawn are presented in the Supplementary Table A.1.

### 2.5 MR data acquisition and preprocessing

Participants took part in a neuroimaging recording session of approximately 2.5 h duration. Before entering the scanner, subjects performed behavioral assessments (see Cognitive assessments), the MR standard safety screening and they were instructed on the details of the session and the current task. In the session, the current learning task and anatomical scan were preceded and followed by a short visual target detection task which is not analyzed here. MRI data were recorded on a Philips Achieva 3 Tesla scanner (Best, The Netherlands) using a 32-element receive head coil. Using a T2-weighted whole-brain gradient-echo planar image sequence, 460 volumes were acquired for each experimental block [Slices = 32; repetition time = 1000 s; echo time = 30 ms; slice gap = 0.5 mm; voxel size = 3 × 3 × 3.5 mm^3^; flip angle = 65°; field of view = 240 × 127.5 × 240 mm^2^; SENSE-factor = 2]. In addition, a field map and a high-resolution T1-weighted anatomical image were acquired.

fMRI data preprocessing and analysis were performed in the SPM12 toolbox. Preprocessing included distortion correction of functional images, slice time correction and coregistration of the functional data to the T1-weighted image. The deformation fields derived from segmentation of the T1 image were used for normalization to the Montreal Neurological Institute (MNI) −152 template space. Last, smoothing with a 6 mm full-width-half-maximum kernel was applied to the functional data. Motion artifacts were assessed by calculating the framewise displacement (FD) values of each subject and task block ^57^. Only subjects with FD < 0.5 mm were included in the analysis (mean 0.18±0.04; no excluded participants based on this criterion). Moreover, single volumes with FD > 1 were censored in the statistical analyses using an additional binary regressor (mean 0.24±0.35 % of volumes excluded; and a maximum of 3.7 % excluded in one participant).

### 2.6 Statistical analysis

#### 2.6.1 Task performance

The basic measures of accuracy (proportion of correct trials) and reaction times (RTs) for correct responses were averaged across thirds of 16 trials (one third of the total number of trials) for each block. Mixed-model ANOVAs were performed on these measures with third (1-3), block (1, 2) and task parts (A, B) as fixed effects and a random intercept. Follow-up analyses examined the effects of the factor third in each task separately. Moreover, associations between task accuracy, RTs, model parameters and cognitive tests were investigated with Pearson correlations and linear regressions. Spearman correlation values were used when data were not normally distributed.

#### 2.6.2 Model-based fMRI analysis

Two GLMs were conducted convolving stimuli and feedback onsets with the hemodynamic response function (as implemented in SPM12) and the different trial-based parameters from the model as parametric modulators. The parameters serving as parametric modulators were AS and PE). The onsets of stimuli and feedback in trials where no responses were given (‘too late’ responses) were added as an additional regressor of no interest. In addition, six realignment parameters from the data preprocessing were included as nuisance regressors and a binary regressor censored scans with FD > 1 (see MR data acquisition and preprocessing).

All fMRI analyses used a cluster-based family-wise error corrected significance threshold of *p*_FWEc_ *=* 0.05 with a cluster-defining threshold of *p*_CDT_ = 0.001.

The whole-brain analysis was further refined by a region of interest (ROI) analysis. ROIs were defined using the meta-analysis tool Neurosynth ^58^. They are summarized in Table 2. The search terms ‘letter’, ‘audiovisual’, ‘learning’ and ‘error’ were used to find the coordinates from areas related to reading, learning and feedback processing areas in Neurosynth. We extracted the first eigenvariate of the time course of active voxels (p < 0.05) within a spherical search volume (*r* = 6 mm) around these coordinates.

**Table 2.** Set of regions of interest based on Neurosynth online meta-analysis search tool.

| keyword/ brain region | MNI coordinates |  |  |
| --- | --- | --- | --- |
|  | x | y | z |
| *‘Letter’* |  |  |  |
| L Fusiform gyrus | -44 | -58 | -14 |
| R Fusiform gyrus<sup>a</sup> | 44 | -58 | -14 |
| L Inferior frontal gyrus<sup>a</sup> | 46 | 2 | 24 |
| R Inferior frontal gyrus | -46 | 2 | 24 |
| *‘Audiovisual’* |  |  |  |
| L Superior temporal gyrus | -52 | -22 | 8 |
| R Superior temporal gyrus | 54 | -24 | 4 |
| *‘Learning’ and ‘error’* |  |  |  |
| L Putamen | -14 | 8 | -10 |
| R Putamen | 26 | 2 | -8 |
| L Hippocampus | -22 | -40 | 4 |
| R Hippocampus | 22 | -32 | 6 |
| L Caudate | -12 | 10 | -10 |
| R Caudate | 14 | 10 | -10 |
| L Insula | -38 | 20 | -6 |
| R Insula | 42 | 18 | -6 |
| L mid Cingulum | 0 | 22 | 38 |
| R mid Cingulum | 4 | 26 | 40 |

L = left hemisphere; R = right hemisphere; <sup>a</sup> Contralateral regions were manually added and not found in the Neurosynth search

Additionally, the appendix presents a conventional fMRI analysis without using RLDDM parameters and dividing the onsets of stimuli and feedback in thirds of 16 trials.

## 3 Results

### 3.1 Behavioural data analyses: Learning performance

#### 3.1.1 Basic analysis of reaction time and accuracy

The main linear mixed models included the factors *part* (A,B), *third* (1,2,3) and *block* (1,2). They were followed by separate models examining learning and potential block effects in each part separately. The descriptive statistics of RT and accuracy (proportion of correct responses) are presented in Table 3 and Figure 3.

**Table 3.** Task accuracy (proportion correct) and reaction times of correct responses.

|  | FBL-A | FBL-B |
| --- | --- | --- |
| | $M(SD)$ | $M(SD)$ |
| Accuracy |  |  |
| Third 1 | 0.61 (0.12) | 0.57 (0.13) |
| Third 2 | 0.80 (0.16) | 0.63 (0.18) |
| Third 3 | 0.85 (0.14) | 0.77 (0.17) |
| Total | 0.75 (0.17) | 0.66 (0.18) |
| RTs |  |  |
| Third 1 | 1365.08 (208.04) | 1529.85 (228.42) |
| Third 2 | 1273.00 (198.13) | 1553.02 (180.03) |
| Third 3 | 1256.97 (191.56) | 1518.75 (207.85) |
| Total | 1298.35 (204.16) | 1533.87 (206.00) |

**Figure 3.**
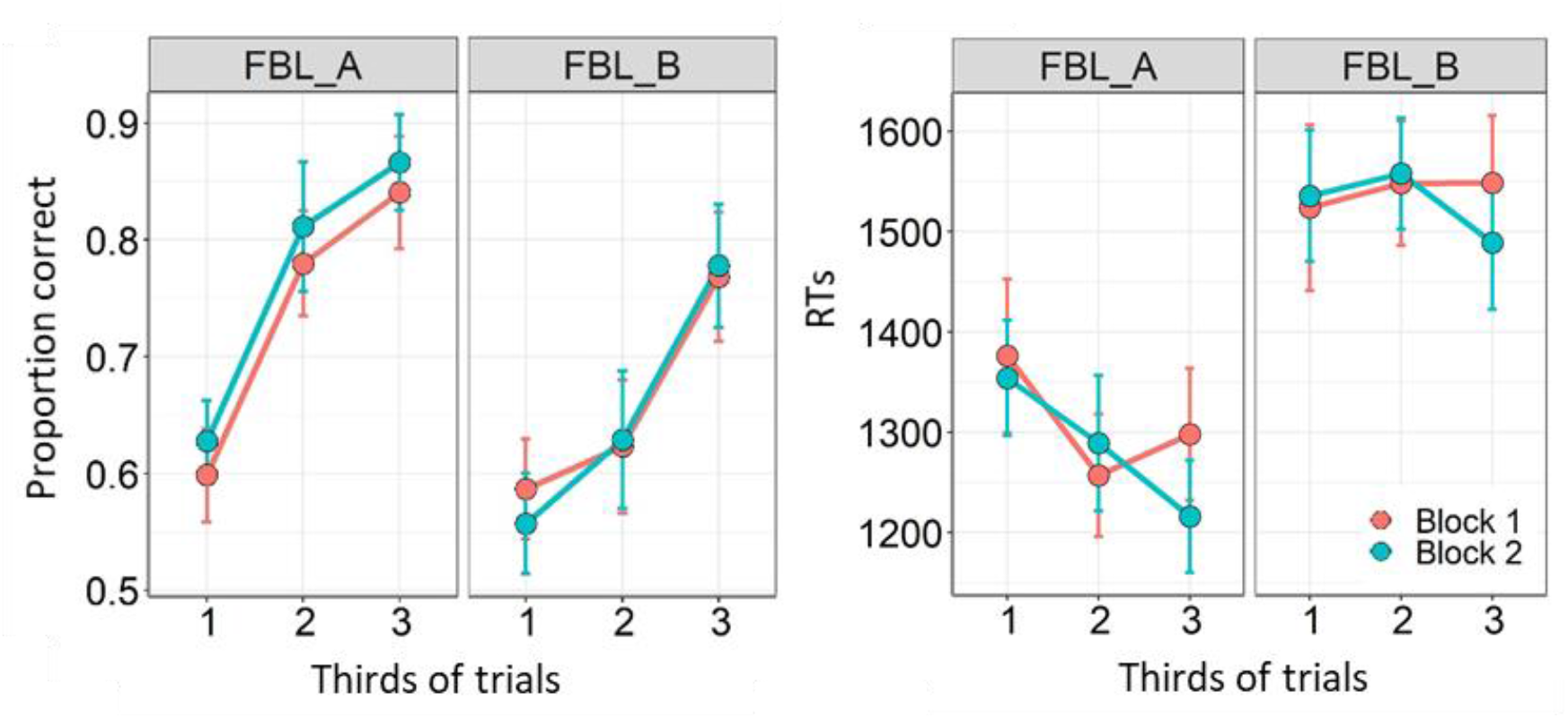
Accuracy (left plots) and RTs (right plots) averaged per trial thirds, separate lines per experimental block (red color indicates first block; blue color the second). Tasks A and B are presented in separate panels.

##### 3.1.1.1 Accuracy

The analysis on the proportion of correct responses yielded a main effect of third (*F*(2,418)= 106.06, *p* < 0.001), indicating the increase in accuracy from the first to the last third of the blocks in both tasks. This learning curve is illustrated in Figure 3 (left panel), see Table 3 for mean accuracy values. Additionally, there was a main effect of part (*F*(1,418)= 61.35, *p* < 0.001) and a significant interaction between part and third, *F*(1,418) =9.37, *p* < 0.001, supporting a different learning curve between parts A and B, as shown in Figure 3. No other effects reached statistical significance in this model (*ps* > 0.176). Further contrasts comparing the part in each third showed higher accuracy in FBL-A vs FBL-B in the third 2 (*t* (418) = 7.89, *p* < 0.001) and third 3 (*t* (418) = 3.74, *p* < 0.001); in third 1 this difference was only detected at trend levels (*p* = 0.055).

The differences in learning rates between FBL-A and FBL-B were further examined in separate models per task part. In both part A and B there was a significant main effect of third, (*F*(2,190) = 75.85, *p* < 0.001) and (*F*(2,190) = 43.58, *p* < 0.001), respectively. No block effects or interaction between thirds and blocks were found significant, although there was a trend at *p* = 0.085 in FBL-A for the main effect of block. The *t* pairwise comparisons between thirds in FBL-A showed significantly increased accuracy over both blocks from third 1 to third 2 (*t* (190) = 8.96, *p* < 0.001), and from third 2 to third 3 (*t* (190) = 2.85, *p* = 0.014). The pairwise comparisons on FBL-B data also showed significant increases from third 1 to third 2 (*t* (190) = 2.43, *p* = 0.042) and from third 2 to third 3 (*t* (190) = 6.59, *p* < 0.001).

To further illustrate these learning curves for each stimulus pair, the Supplementary Figure A.3 shows the cumulative summed probabilities for each block and task.

##### 3.1.1.2 Reaction times

The linear mixed model on the reaction times of correct responses revealed a main effect of third (*F*(2,418) = 6.46, *p* = 0.002) showing shorter RTs in the later trials of the blocks. The mean RTs are shown in Table 3 and Figure 3 (right panel). In addition, there was a main effect of part (*F*(1,418)= 300.32, *p* < 0.001) showing larger RTs in FBL-B compared to FBL-A (see histograms in Supplementary Figure A.4.). The part effect was also found in interaction with third, (*F*(1,418)= 6.92, *p =* 0.001). In addition there was a significant interactions between third and block (*F*(2,418)= 4.03, *p =* 0.018), suggesting more pronounced effects of third in block 2 (follow up pairwise comparison only showed significance in the third 2 vs third 3 comparison in block 2; *t*(417) = 2.74, *p* = 0.017). No other effects were significant in the main model, *ps >* 0.179.

The separate analyses per task part followed the main effect of part and the interaction between part and third. The analysis of FBL-A revealed a main effect of third (*F* (2,190) = 12.66, *p* < 0.001), suggesting shorter RTs as the block progressed (pairwise comparisons: third 1 vs third 2, *t*(190) = −3.97, *p* < 0.001; third 2 vs third 3 not significant, *p* = 0.769). There was also a trend for an interaction between third and block (*F* (2,190) = 3.03, *p =* 0.050) suggesting that this effect was more pronounced in block 2. The analysis on FBL-B revealed no significant effects or interactions on the mean RTs.

To sum up the basic performance analysis, we found FBL-B compared to FBL-A yielded longer RTs and lower accuracy. Both parts showed pronounced learning curves, i.e., increased accuracy from first to last third of trials, but the increase was somewhat delayed in the FBL-B, and the accuracy differences between the tasks were mainly found in the second and last third.

#### 3.1.2 Reinforcement-learning drift-diffusion model

The model with dual learning showed a better fit for our data in the model comparisons (see Methods and supplementary Table A.2). The posterior distributions of the subject-level parameters derived from the model are presented in Figure 4. Scatter plots and distribution of these parameters are shown in Supplementary Figure A.5.

**Figure 4.**
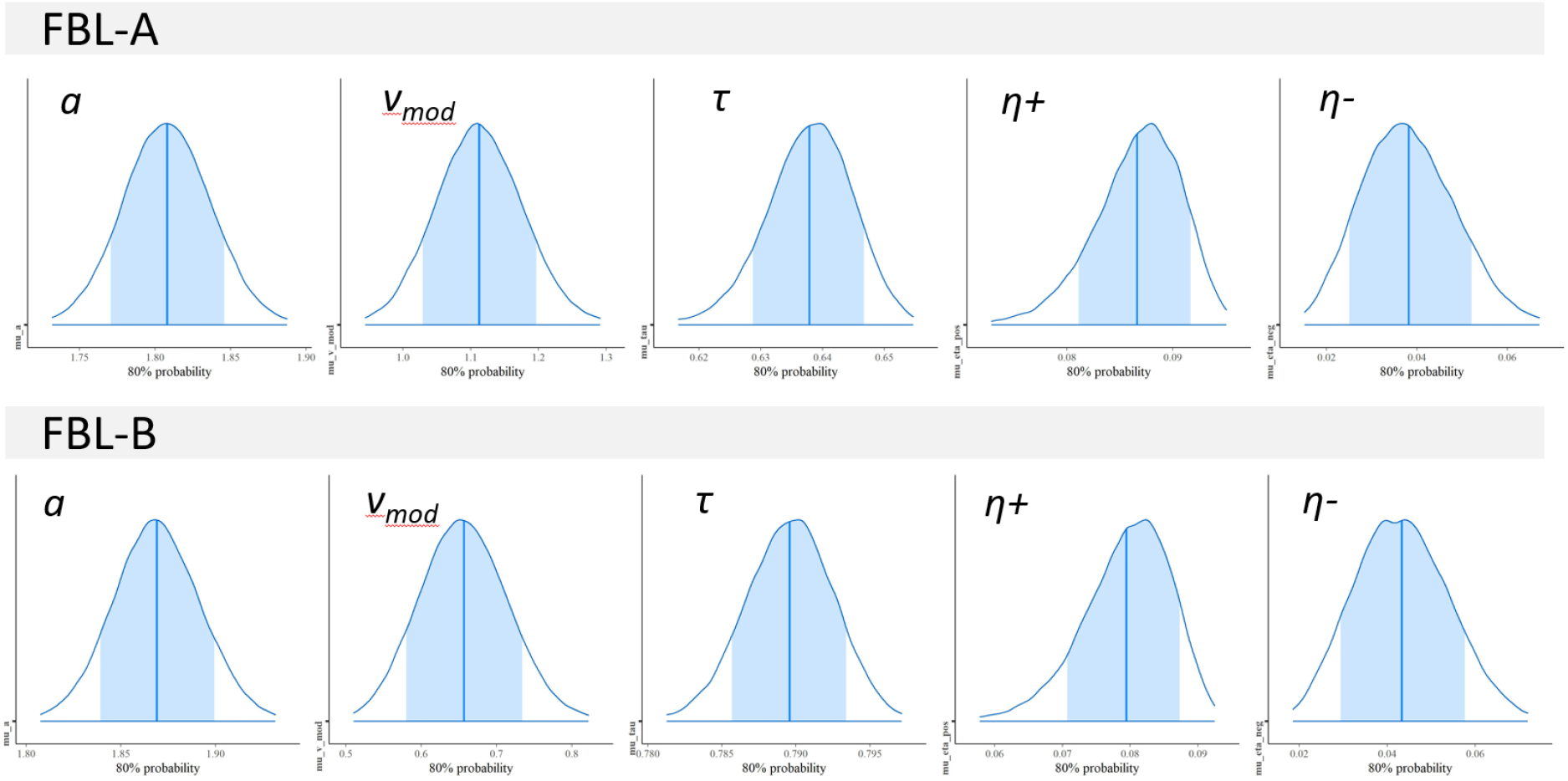
Posterior distributions of the subject-level parameters for each task. Shaded areas represent 80% credibility intervals, vertical line represents the mean point estimate.

#### 3.1.3 Correlations between task performance and cognitive skills

As an additional analysis we explored the correlations between performance in our experimental tasks (captured by RTs, accuracy and model parameters) and cognitive abilities. This secondary analysis is reported in the Appendix (see Supplementary Table A.3). The analysis yielded potentially interesting associations indicating better task performance with higher non-verbal IQ, RAN and text reading scores.

### 3.2 Changes in neural activations with learning

#### 3.2.1 Model-based fMRI

The goal of the main analysis in this study was to identify patterns of activation associated with trial-by-trial learning and prediction error. In the first-level analysis the RLDDM parameters association strength (AS) and prediction error (PE) were mean centered and entered as parametric modulators of stimuli and feedback onsets, respectively, in the GLMs convolving the onsets with the hemodynamic response function.

##### 3.2.1.1 Model-based whole-brain

The tables 4.a. and 4.b show suprathreshold clusters for adjusted *p_FWE_* < 0.05. The main activation clusters are also illustrated in Figure 5. The results show clusters in the visual cortex positively modulated by AS in both parts. A more extensive set of areas showed positive modulation by PE, including areas in the orbitofrontal cortex, striatal regions of putamen, caudate and hippocampus as well as occipital and temporal regions, and a large cluster in the postcentral region. Both parts showed similar patterns although the extent of activations is more widespread in part A. Finally, there was a negative modulation by prediction error in the inferior frontal region, including the insula, and in part B in the supplementary motor area. The latter result was only detected in few voxels (k = 16) in part A (not reported in table).

**Table 4.a.** Results from model-based fMRI in FBL-A.

| Contrast | Brain area | MNI coordinates | | | cluster<br>$p_{FWE_{cor}}$ | Cluster<br>k | Peak<br>Z | Peak<br>T | peak<br>$p_{FWE_{cor}}$ |
| --- | --- | --- | --- | --- | --- | --- | --- | --- | --- |
|  |  | X | Y | Z |  |  |  |  |  |
| *Stimulus onset* |  |  |  |  |  |  |  |  |  |
| AS+ | Calcarine L | -11 | -87 | -3 | < .0001 | 112 | 6.46 | 8.81 | < .0001 |
|  | Lingual R | 16 | -72 | -12 | < .0001 | 39 | 5.86 | 7.55 | 0.0002 |
| *Feedback onset* |  |  |  |  |  |  |  |  |  |
| PE+ | Angular L | -50 | -72 | 27 | < .0001 | 125 | 7.08 | 10.32 | < .0001 |
|  | Postcentral R | 31 | -33 | 48 | < .0001 | 1865 | 6.77 | 9.52 | < .0001 |
|  | Frontal Med Orb L | -8 | 42 | -12 | < .0001 | 568 | 6.72 | 9.41 | < .0001 |
|  | Putamen L | -14 | 9 | -9 | < .0001 | 60 | 6.54 | 8.98 | < .0001 |
|  | Sup Frontal L | -17 | 27 | 48 | < .0001 | 78 | 6.39 | 8.63 | < .0001 |
|  | Mid Frontal Orb L | -29 | 42 | -12 | < .0001 | 59 | 6.31 | 8.46 | < .0001 |
|  | ParaHippocampal L | -17 | -3 | -27 | < .0001 | 67 | 6.12 | 8.06 | < .0001 |
|  | Precuneus R | 7 | -54 | 18 | < .0001 | 262 | 6.05 | 7.92 | < .0001 |
|  | Calcarine R | 28 | -63 | 18 | < .0001 | 44 | 5.83 | 7.48 | 0.0002 |
|  | Lingual L | -26 | -45 | 3 | < .0001 | 35 | 5.7 | 7.23 | 0.0004 |
|  | Sup Temporal R | 70 | -3 | 6 | < .0001 | 58 | 5.66 | 7.16 | 0.0006 |
|  | Caudate R | 19 | 3 | 24 | < .0001 | 59 | 5.51 | 6.89 | 0.0013 |
|  | Caudate L | -23 | -3 | 21 | < .0001 | 30 | 5.42 | 6.72 | 0.0022 |
|  | ParaHippocampal R | 25 | -24 | -21 | < .0001 | 30 | 5.38 | 6.65 | 0.0027 |
| PE- | Inf Orbito Frontal R | 37 | 24 | -9 | < .0001 | 115 | 6.42 | 8.72 | < .0001 |

A voxel-wise threshold of $p_{FWE_{corr}} < 0.05$ was used and only clusters with $k > 30$ are reported. Mid = middle; Sup = superior; Inf = inferior; Med = medial; Orb= orbital.

**Table 4.b.**
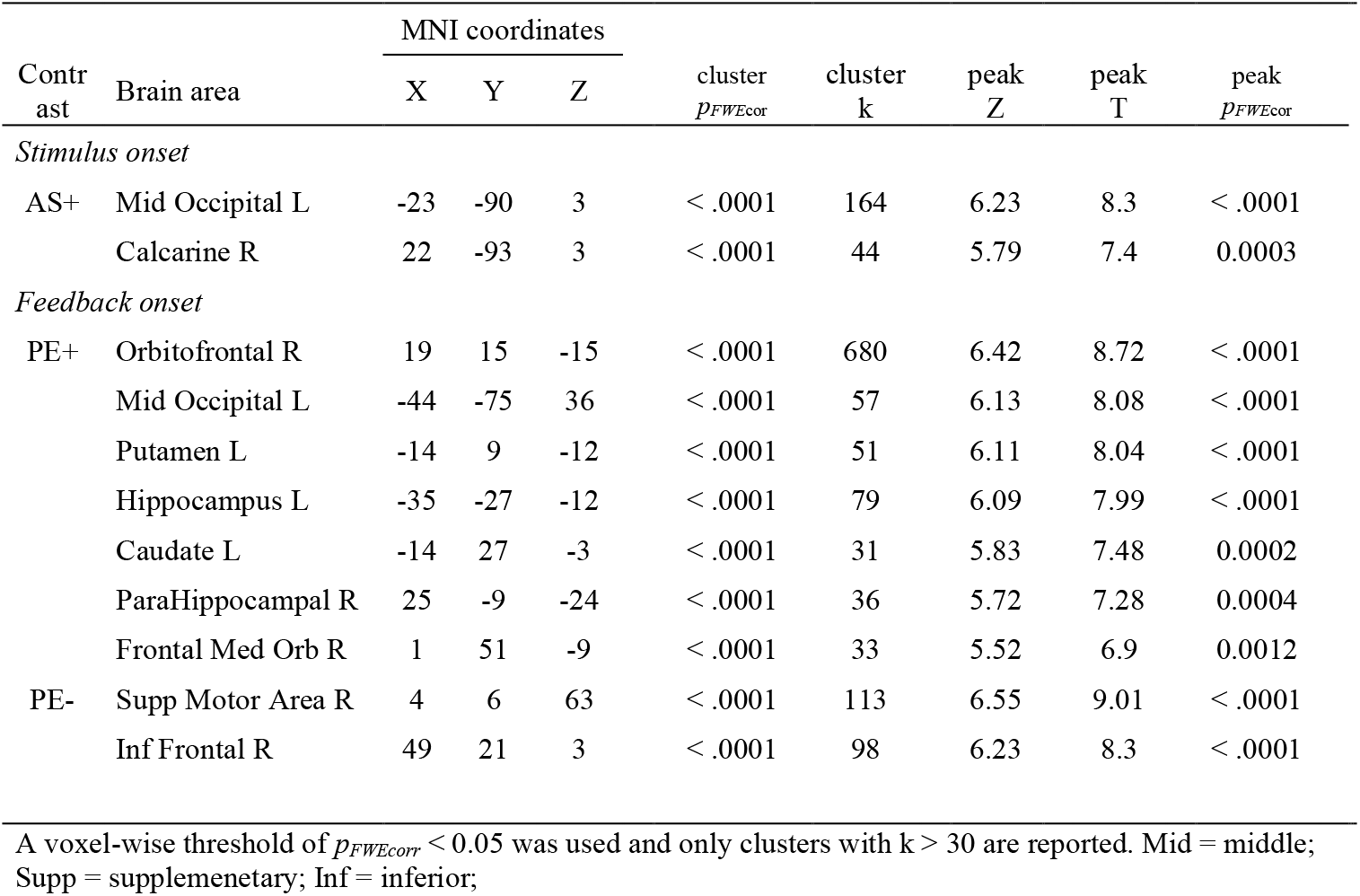
Results from model-based fMRI in FBL-B.

| Contrast | Brain area | MNI coordinates |  |  | cluster<br>*p*<sub>FWEcor</sub> | cluster<br>k | peak<br>Z | peak<br>T | peak<br>*p*<sub>FWEcor</sub> |
| --- | --- | --- | --- | --- | --- | --- | --- | --- | --- |
|  |  | X | Y | Z |  |  |  |  |  |
| *Stimulus onset* |  |  |  |  |  |  |  |  |  |
| AS+ | Mid Occipital L | -23 | -90 | 3 | < .0001 | 164 | 6.23 | 8.3 | < .0001 |
|  | Calcarine R | 22 | -93 | 3 | < .0001 | 44 | 5.79 | 7.4 | 0.0003 |
| *Feedback onset* |  |  |  |  |  |  |  |  |  |
| PE+ | Orbitofrontal R | 19 | 15 | -15 | < .0001 | 680 | 6.42 | 8.72 | < .0001 |
|  | Mid Occipital L | -44 | -75 | 36 | < .0001 | 57 | 6.13 | 8.08 | < .0001 |
|  | Putamen L | -14 | 9 | -12 | < .0001 | 51 | 6.11 | 8.04 | < .0001 |
|  | Hippocampus L | -35 | -27 | -12 | < .0001 | 79 | 6.09 | 7.99 | < .0001 |
|  | Caudate L | -14 | 27 | -3 | < .0001 | 31 | 5.83 | 7.48 | 0.0002 |
|  | ParaHippocampal R | 25 | -9 | -24 | < .0001 | 36 | 5.72 | 7.28 | 0.0004 |
|  | Frontal Med Orb R | 1 | 51 | -9 | < .0001 | 33 | 5.52 | 6.9 | 0.0012 |
| PE- | Supp Motor Area R | 4 | 6 | 63 | < .0001 | 113 | 6.55 | 9.01 | < .0001 |
|  | Inf Frontal R | 49 | 21 | 3 | < .0001 | 98 | 6.23 | 8.3 | < .0001 |

A voxel-wise threshold of $p_{FWEcor} < 0.05$ was used and only clusters with $k > 30$ are reported. Mid = middle; Supp = supplementary; Inf = inferior;

**Figure 5.**
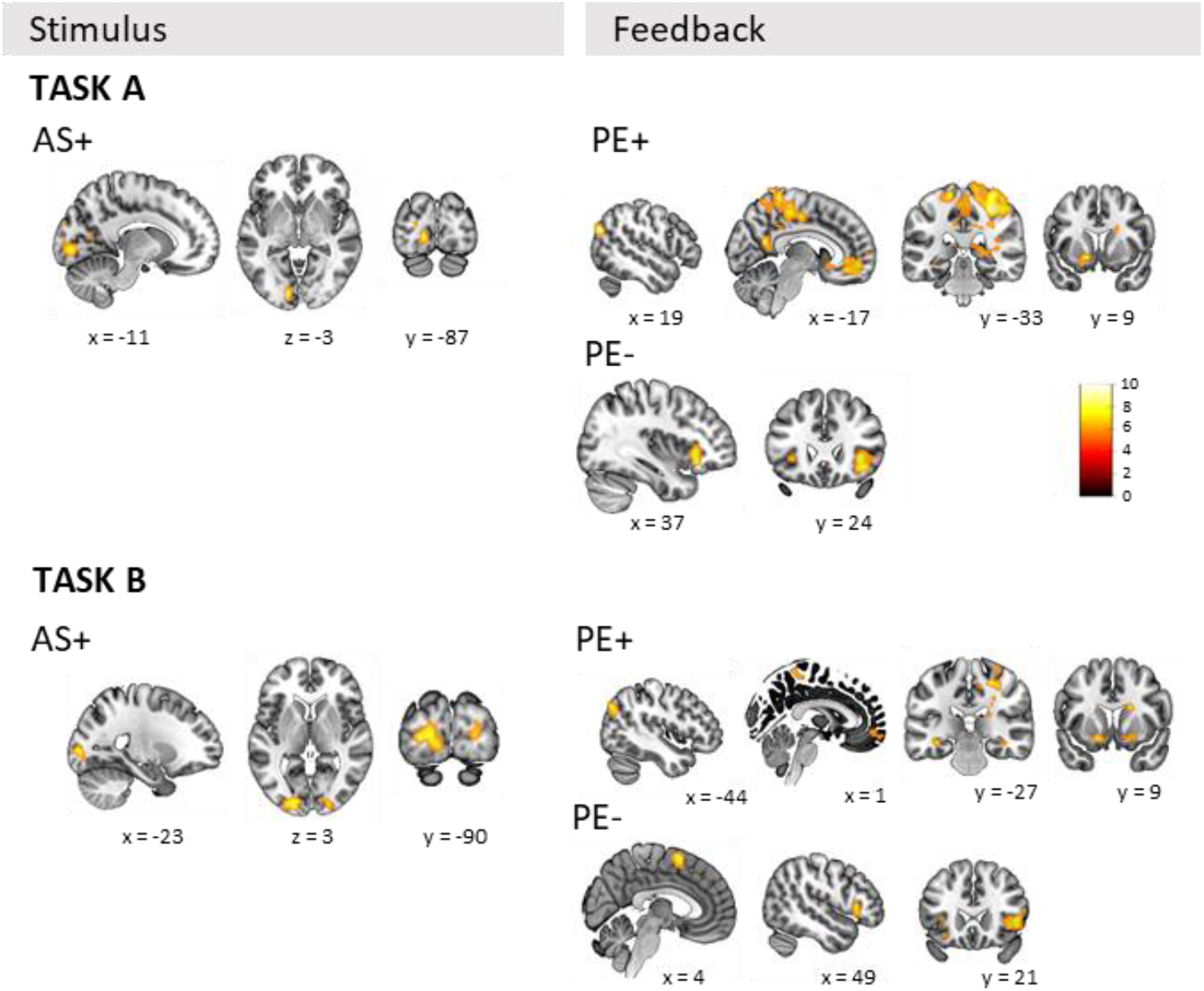
Suprathreshold clusters from the model-based analysis at *p_FWE_*_cor_ < 0.05. Trial-level parameters from the computational model were used as parametric modulators. AS+ = positive influence of associative strength in processing stimuli; PE+ = positive influence of the prediction error in processing feedback. Coordinates in MNI-152 space.

##### 3.2.1.2 Model-based ROI

The ROI analysis is presented in Table 5 and Figure 6. The results showed a positive modulation of activation during stimulus processing by AS in the left fusiform and left superior temporal gyrus in both parts. An additional effect in the left inferior frontal gyrus was found in part B. Regarding PE and feedback processing, there was positive modulation in bilateral putamen, caudate and hippocampus in both task parts, and an additional effect in bilateral superior temporal gyrus was found in part A. In the opposite direction, insula and cingulum were negatively associated with PE.

**Table 5.** ROI t-tests against zero in model-based effects AS+ and PE+.

| keyword/ brain region | FBL-A |  |  |  | FBL-B |  |  |  |
| --- | --- | --- | --- | --- | --- | --- | --- | --- |
|  | Stimuli AS+ |  | Feedback PE+ |  | Stimuli AS+ |  | Feedback PE+ |  |
|  | *t* | *p<sub>FDR</sub>* | *t* | *p<sub>FDR</sub>* | *t* | *p<sub>FDR</sub>* | *t* | *p<sub>FDR</sub>* |
| L Fusiform | **3.62** | **0.003** | 1.44 | 0.231 | **2.66** | **0.032** | 0.53 | 0.788 |
| R Fusiform | 0.88 | 0.43 | 2.17 | 0.079 | 1.62 | 0.215 | 0.67 | 0.768 |
| L IFG | 1.89 | 0.117 | 0.79 | 0.464 | **2.56** | **0.037** | -1.03 | 0.491 |
| R IFG | 1.74 | 0.144 | -1.2 | 0.316 | -0.27 | 0.866 | -2.27 | 0.062 |
| L STG | **3.04** | **0.011** | **2.43** | **0.046** | **2.58** | **0.037** | 0.51 | 0.788 |
| R STG | 1.92 | 0.117 | **4.27** | **<.001** | **2.51** | **0.039** | 1.88 | 0.136 |
| L Putamen | 1.95 | 0.117 | **7.16** | **<.001** | -0.01 | 0.995 | **6.53** | **<.001** |
| R Putamen | 1.45 | 0.231 | **4.45** | **<.001** | 1.18 | 0.424 | **6.07** | **<.001** |
| L Hippocampus | -1.08 | 0.34 | **5.25** | **<.001** | -0.38 | 0.866 | **5.45** | **<.001** |
| R Hippocampus | -0.34 | 0.733 | **5.71** | **<.001** | 0.22 | 0.88 | **4.98** | **<.001** |
| L Caudate | 1.17 | 0.316 | **8.28** | **<.001** | 0.6 | 0.788 | **9.29** | **<.001** |
| R Caudate | 1.79 | 0.138 | **6.38** | **<.001** | -1.16 | 0.424 | **7.34** | **<.001** |
| L Insula | 1.3 | 0.280 | **-4.76** | **<.001** | -0.27 | 0.866 | **-5.65** | **<.001** |
| R Insula | 1.15 | 0.316 | **-6.83** | **<.001** | 0.17 | 0.891 | **-5.82** | **<.001** |
| L mid Cingulum | 0.87 | 0.43 | **-2.8** | **0.02** | 0.51 | 0.788 | **-4.14** | **<.001** |
| R mid Cingulum | -0.68 | 0.519 | **-3.97** | **<.001** | 0.33 | 0.866 | **-5.67** | **<.001** |

mid cingulum = median cingulate gyrus; left hemisphere; R = right hemisphere; STG= superior temporal gyrus; IFG = inferior frontal gyrus; AS+ = positive influence of association strength on stimuli processing; PE+ = positive influence of prediction error on feedback processing; Bold text indicates results with *p<sub>FDR</sub>* < 0.05 (False Discovery Rate adjustment for 32 tests).

**Figure 6.**
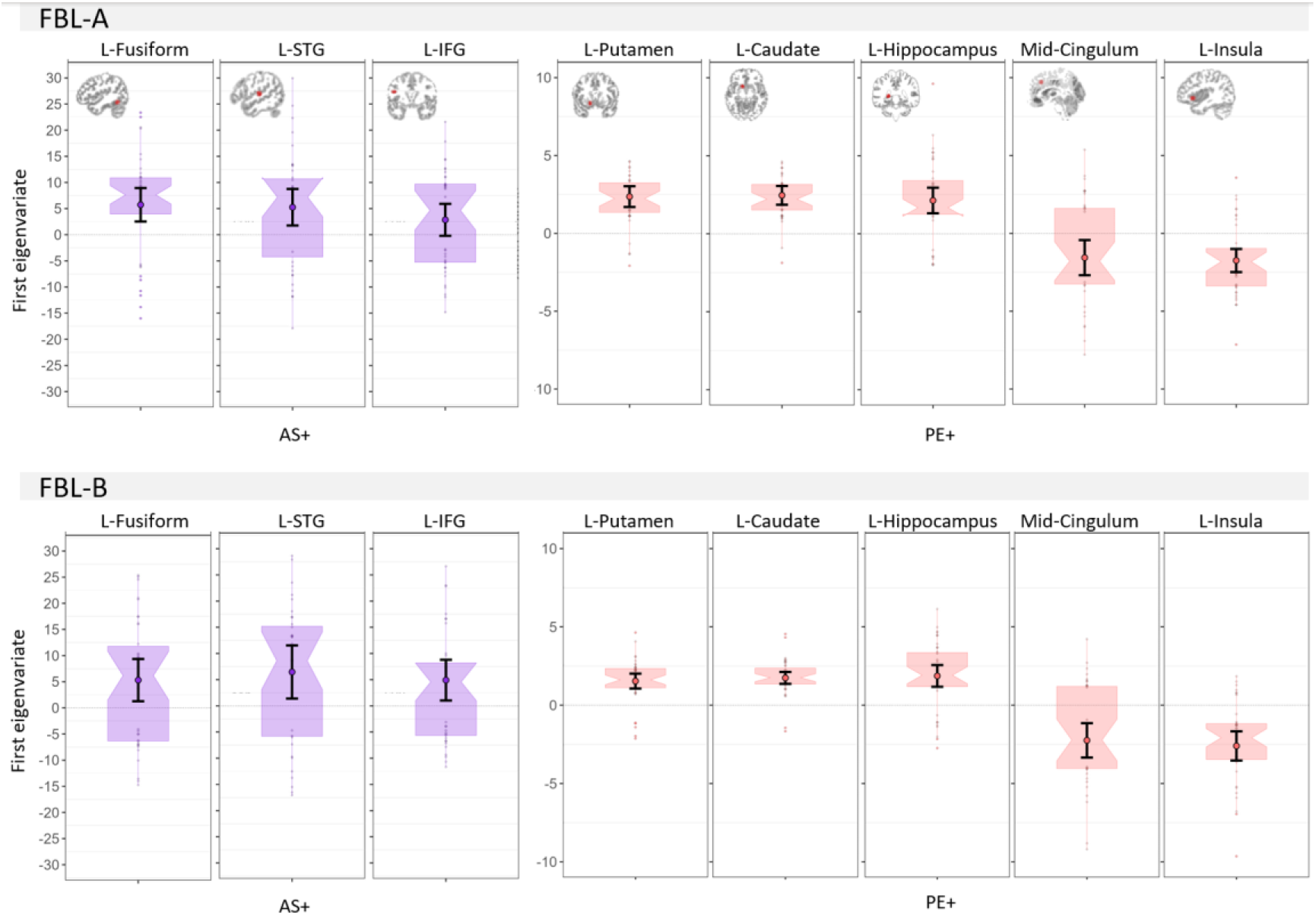
First eigenvariates for the region of interest (ROI) showing significant modulations by associative strength (AS) for processing stimuli and by prediction error (PE) on feedback presentations. Only left hemisphere ROIs are shown. L=left; STG=superior temporal gyrus; IFG=inferior frontal gyrus. Boxplots are notched around the median and error bars represent 95 % CI. The embedded brain images show the corresponding ROI mask.

## 4 Discussion

The main goal of this study was to characterize the brain systems implicated in learning new symbol and speech sound associations and to assess individual differences in associative learning. For this, we used two task parts in which symbol-speech sound associations were learned via feedback on screen. As expected, we found that associations in part A were easier to learn than the ones of the part B, in which diacritic marks were added to modify previously learned sounds. A reinforcement learning drift diffusion model was used to model performance in each part and the trial-by-trial parameters of association strength and prediction error were used as parametric modulators of stimulus and feedback processing, respectively, in the model-based fMRI analysis. The analysis yielded similar patterns of activations in both parts, showing how audiovisual associative learning contributes to visual specialization, the role of striatal regions in prediction error encoding and the progressive engagement of cingulate and frontal regions related to performance monitoring.

### 4.1 Occipital responses to stimuli increase with strengthening of audiovisual associations

One of our goals was to examine specialization in visual/audiovisual regions when learning new associations within the short time-scale of an experimental task. The model-based analysis revealed an occipital region that became more active for processing stimuli with increasing association strength. This region was located in the medial posterior portion of the occipital cortex with a peak in the left hemisphere (see tables 3.a and b). The peak location approximates the letter-selective region selection described in a study using intracerebral recordings ^59^. In that study, the electrode contacts that responded selectively to letters, were widely distributed across regions of the ventral occipital cortex (e.g., inferior occipital, medial fusiform, middle temporal and inferior temporal gyrus and other neighboring regions). The highest proportion of letter-selective intracerebral electrodes were located in the (left) posterior portion of the inferior occipital cortex, while a smaller, more anterior located group of electrodes detected selective responses in prelexical and lexical contrasts. A similar posterior-anterior gradient for letter-word selectivity was found in a previous fMRI study, which referred to a putative a ‘letter-form area’ in the posterior fusiform gyrus ^60^. The peak of the cluster modulated by association strength in our model-based analysis has a more posterior and medial location similar to previous reports ^59,60^. Those studies also detected more anterior letter-selective regions, interpreted as letter form areas in their tasks, which included letter strings. Our ROI analysis also suggested that the activation extended to the left fusiform region.

Several factors may contribute to the posterior location in our results and the previously reported ‘letter form area’. The current study presents single characters instead of strings. In contrast to our relatively simple stimuli, more complex visual items could be expected to elicit stronger activations and possibly recruit a more extensive visual region. For example, a study comparing consonant-vowel-consonant strings to single letters yielded more activation in occipital regions to the string stimuli ^61^. Moreover, we used false fonts learned over a short experiment instead of real letters, and our task did not require semantic processing, as it was required in ^60^ and which may explain the extension of their letter selective response also to more anterior vOT areas. In relation to this, the left posterior middle temporal gyrus has been associated with lexical/semantic processing by previous work ^62,63^. In addition, a previous functional and structural MRI study, found an area around the middle occipito-temporal sulcus that was sensitive to linguistic information, i.e., contrasts involving real vs pseudowords and falsefonts ^64^. Of note, the coordinates of the occipital region modulated by association strength in the current study was similar in both parts of the task, even though part B may require more focus on detailed features (diacritic marks). This would suggest that processing of visual features is not driving this result, although this would require further examination in a visual experiment with direct comparison between stimuli. Importantly, the modulation by the association strength parameter points at audiovisual learning, rather than just growing visual familiarity during the block, as a main contributor to the increase in visual activations.

To sum up, our model-based fMRI result supports results from developmental studies showing facilitated visual processing after audiovisual integration when learning how to read ^11,16,65–67^. Also in line with this, but related to broader sensory learning and conditioning, top-down influences from auditory to visual cortex were found in a target detection task in which unknown to participants, auditory distractors predicted the presence or absence of visual stimuli^68^.

Besides these findings, we should note that the supplementary, conventional analysis (see Supplementary Figure A.6 and tables A.4.a and b) using thirds of trials, confirmed stronger activations in extended areas of occipital and temporal regions in the consolidation phase compared to the initial trials of the block in both tasks. These effects may reflect enhanced representations after crossmodal learning, similar to those found with reading acquisition ^2,11,16,69^. We should note that from these analyses we cannot rule out the influence of mere repeated exposure, although previous studies suggest repetition suppression effects rather than stronger activations to repeating, non-degraded visual stimuli ^70^. On the other hand, building up capacity to perceptually discriminate the symbols, i.e., familiarity, could also be an alternative modulator of activations in the last part of the task ^23^. In addition, the whole-brain conventional analysis also showed inferior frontal and striatal regions, like caudate and putamen, that were more active in the last part of the block. This suggests engagement of structures import for consolidating learning. Interestingly, the putamen was previously related to anticipation of outcomes from stimuli in a stimuli-action-reward learning task ^71^. Thus, these results may reflect stronger capacity to select the correct response as participants learnt the false font-speech sound associations.

### 4.2 Striatal regions involved in prediction error for audiovisual learning

Another goal of the current study was to investigate the learning mechanisms and brain regions contributing to visual/audiovisual specialization. Prediction error (PE) is proposed to drive learning by signaling the need to update predictions to minimize the discrepancy between expectations and a given outcome ^72^. Thus, average PE decreases as predictions become more accurate and learning progresses ^29^. In the current setting, PE in each trial determines the change in association strength.

In both task parts, PE-dependent activations in several subcortical regions including putamen, caudate and hippocampal region, as well as posterior occipital cortex and medial frontal cortex. In the ROI analysis, putamen, hippocampus and caudate activity varied with the model-derived PE in both parts of the task. This finding supports the notion that striatal nuclei are involved in encoding PE signals ^31,32^. Besides sharing a common role in skill learning and memory ^73^, the nuclei in the striatum have been associated with different connections and functions. For example, evidence suggests that caudate is functionally connected with frontal regions ^74,75^, and with orbitofrontal and cingular cortices ^76^. On the other hand, the putamen has more basic sensorimotor connections ^74,77^. From the functional viewpoint, generally, anterior putamen, caudate and their connections to association cortices are considered to be part of a broader cognitive system, while ventral striatal areas form a reward system and posterior putamen is more associated with a motor system ^76^. It should be noted that the anatomical and functional boundaries between striatal systems are not always clear and, since PE is considered a rather ubiquitous mechanism that works at multiple levels, PE signals have been reported in many different regions depending on whether the focus is on motivational, attentional, cognitive or perceptual responses (see review in ^78^). Close to the associative learning context of the current study, Den Ouden and colleagues (2009) found putamen and primary visual cortex to reflect the magnitude of surprise, i.e., unsigned prediction errors, in their audiovisual task with conditioning ^68^.

The additional activations in the hippocampal region in the current study could also relate to the memory demands of the current task, especially in the learning phase when participants need to keep track of the different visual stimuli, the presented speech sound, as well as their response and subsequent feedback. Previous work suggested a role of hippocampus in selective attention ^79^ and in supporting working memory, especially when binding high resolution cross modal information and handling complex features ^80^. Thus, the hippocampus would be relevant when considering the multisensory aspects of working memory ^80^ and the audiovisual nature of our task. In addition, an interaction between hippocampus and medial prefrontal cortex, both of which showed a positive association with PE in the current study, has also been suggested as key to memory formation and consolidation ^81^.

### 4.3 Prefrontal and cingulate regions negatively associated with prediction error

In both tasks the ROIs insula and mid cingulum, showed a negative association with PE (see table 5). The insula’s posterior regions connect to sensory regions and association cortex, while the anterior region connect to anterior cingulate cortex, ventromedial prefrontal cortex, amygdala and ventral striatum ^82^. The anterior insula function has been associated with error awareness ^83^, certainty and risk evaluation ^84^, anticipation of outcomes ^85^ and risk-averse behavior ^86^, as well as to integrate interoceptive signals with emotional, cognitive and motivational signals ^82^. Together with the anterior cingulate cortex, the anterior insula is proposed to be involved in evaluation of events/stimuli salience and oriented responses (see review in ^87^). The cingulate cortex, generally associated with conflict monitoring and feedback evaluation ^27^, is proposed to facilitate response selection through its connections with mid-cingulate and supplementary motor area ^88^. In addition, the whole-brain analysis, the cluster showing a negative influence by PE, also included the adjacent supplementary motor area (suprathreshold voxels under FWE correction were detected in both tasks, although results in part A were not considered due to the small number of voxels, see tables 4.a and b). This region was also previously linked to learning stimulus-response sequences, in combination with striatum and middle frontal gyrus ^89^.

### 4.4 Limitations

There are a few limitations worth mentioning in the current study design. The additional part B was incorporated in our paradigm to increase difficulty and simulate inconsistent symbol-speech sound associations with diacritic marks as speech sound modifiers. This resulted in the expected difference in performance, suggesting part B to be more demanding. However, a direct comparison of neural activations would be confounded by several factors. The visual load of each task differs as false fonts in part A consist of single characters while part B requires identifying the main character and the modifier mark on top. In addition, the main false fonts in B are not novel as they had been learned in the preceding block from part A. Our fMRI results in each task suggest similar patterns, although considering together task requirement and brain results, part A may provide with a framework that is less ambiguous for interpretation. Finally, the current study is limited in its capacity to relate task performance to relevant reading skills. The current sample presents variability in cognitive performance, however, most subjects performed within normal skill range. Inclusion of poor or dyslexic readers, and a developmental population would allow further assessment of the clinical potential of this paradigm, given the importance of early failure to establish letter-speech sound associations in reading disorders. The correlational analysis between cognitive skills and task performance and model parameters (see Appendix 8.8) supports the potential interest of such investigation.

### 4.5 General conclusions

To conclude, the current results directly relate fast specialization of visual areas with learning symbol-speech sounds associations. This finding is relevant to confirm hypotheses derived from developmental and training studies focusing on broader time windows. In addition, we identified a set of striatal, cingulate, and prefrontal regions engaged in processing feedback and prediction errors when learning these associations. This is an important contribution to reading research as studies focusing on prediction errors in this particular context were largely lacking. The current approach would be suited for further investigations with clinical and developmental samples. In sum, we provide a framework to advance our models of the neurobiology of symbol/letter-speech sound binding, a crucial step when learning to read alphabetic orthographies.

## 5 Acknowledgments

We would like to express our gratitude to all participants in the study.

## 6 Funding

University of Zurich, UZH Postdoc Grant, grant no. [FK-19-040] to GFG. NCCR Evolving Language (SNSF 1NF40_180888)

### 7 Data availability

Data will be made available upon written request to the corresponding author. A formal data sharing agreement will be required. The code is available in the public repository: https://github.com/gorkafraga/GRAPHOLEMO

