## Appendix for "Neural representation of association strength and prediction error during novel symbol-speech sounds learning"

### *Task design*

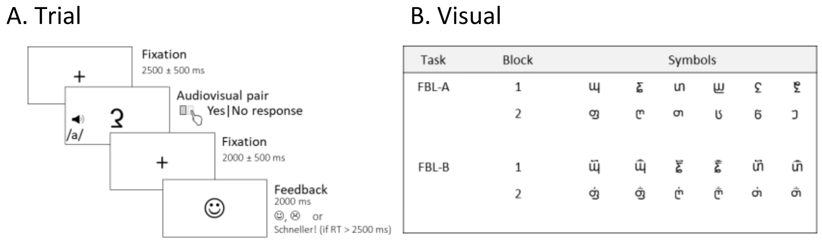

Figure A.1.Task design. (A) Trial design in which a symbol and a phoneme are presented simultaneously and response terminated. Feedback is presented for 2000 ms and indicates whether the response is correct, incorrect or too slow. (B) Table showing the visual stimuli presented in each part and block.

### *RLDDM parameters*

Supplementary Table A.1. Model parameters

| Parameter | Symbol | Value |
|---|---|---|
| Group-level priors | | |
| Mean | $\mu_d$ | ~ $N$ (0,1) |
| Standard deviation | $\sigma_d$ | ~ $N$ (0,0.2) |
| Learning rates means | $\mu_{\eta+/-}$ | ~ $N$ (0,0.3) |
| Learning rates std. deviation | $\sigma_{\eta+/-}$ | ~ $N$ (0,0.5) |
| Subject-level priors | | |
| Learning rates | $\eta_{+/-}$ | ~ $N$ (0,1.5) |
| Decision boundary | $a$ | ~ $exp\ (N(\mu_a, \sigma_a))$ |
| Drift rate modulator | $v_{mod}$ | ~ $exp\ (N(\mu_{v_{mod}}, \sigma_{v_{mod}}))$ |
| Non-decision time | $\tau$ | ~ $\mid \Phi(N(\mu_\tau, \sigma_\tau)) \cdot (RT_{min} - rb)$ + rb $\mid$ |
| Positive learning rate | $\eta_+$ | ~ $0.1 \cdot \Phi(N(\mu_{\eta+}, \sigma_{\eta+})$ |
| Negative learning rate | $\eta_-$ | ~ $0.1 \cdot \Phi(N(\mu_{\eta-}, \sigma_{\eta-})$ |

*Model diagnostics*

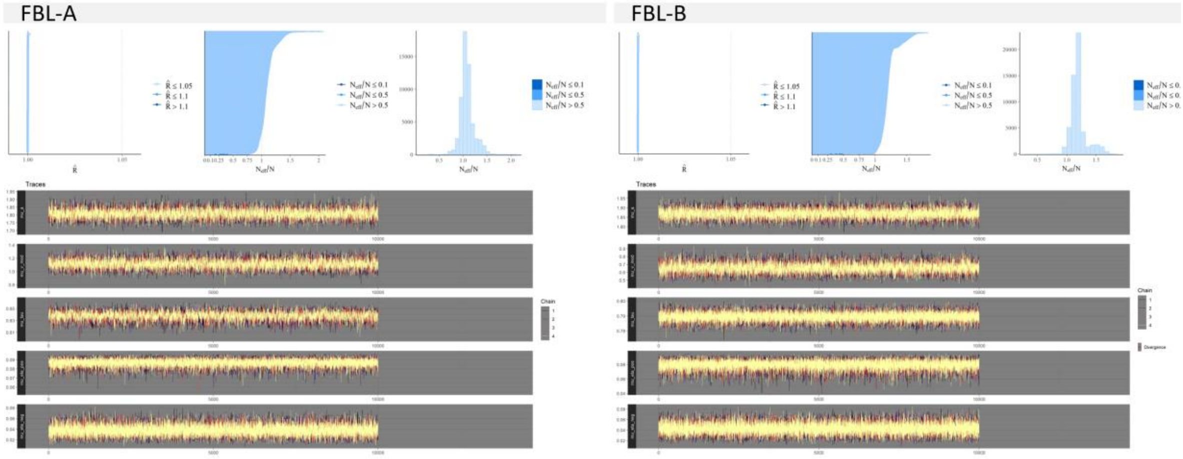

Figure A.2. Model diagnostics for the RLDDM-2 (model with dual learning rates) with 4 chains of 10000 iterations (+ 4000 burn-in) for parts A and B. Top panels $\hat{R}$ convergence estimate (left) and ratios of effective sample size to total sample size (middle and right). Bottom plots are trace plots of the four chains for the group parameters.

*Task accuracy: cumulative probabilities for each stimuli*

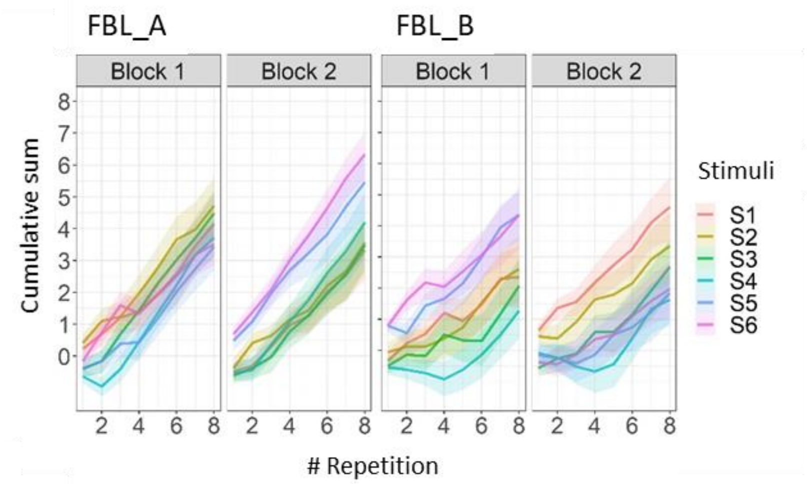

Figure A.3. Learning trajectories per block and part for each unique sound. The values used in the calculation of the cumulative sum are +1 for trials with correct responses, -1 for incorrect trials and 0 for trials with missing responses. Each sound is presented 8 times.

### *Task reaction times: histograms per part and response type*

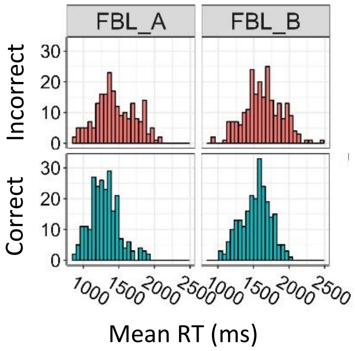

Figure A.4.Histograms of reaction times over all blocks for correct (blue) and incorrect (red) responses. The left panel shows the data for part A and the right panel the data for part B.

### *Table of model comparisons*

Table A.2. Model variation and comparison of fit estimates

| Model | $\eta$ | **FBL A** | | | **FBL B** | | |
|---|---|---|---|---|---|---|---|
| | | *WAIC* | *-elpd* | *Rank* | *WAIC* | *-elpd* | *Rank* |
| RLDDM 1 | Single | 7017.29 | 3508.645 | 2 | 8491.748 | 4245.874 | 2 |
| RLDDM 2 | Dual | 7003.93 | 3501.965 | 1 | 8484.534 | 4242.267 | 1 |

### *Distribution of subject-level model parameters*

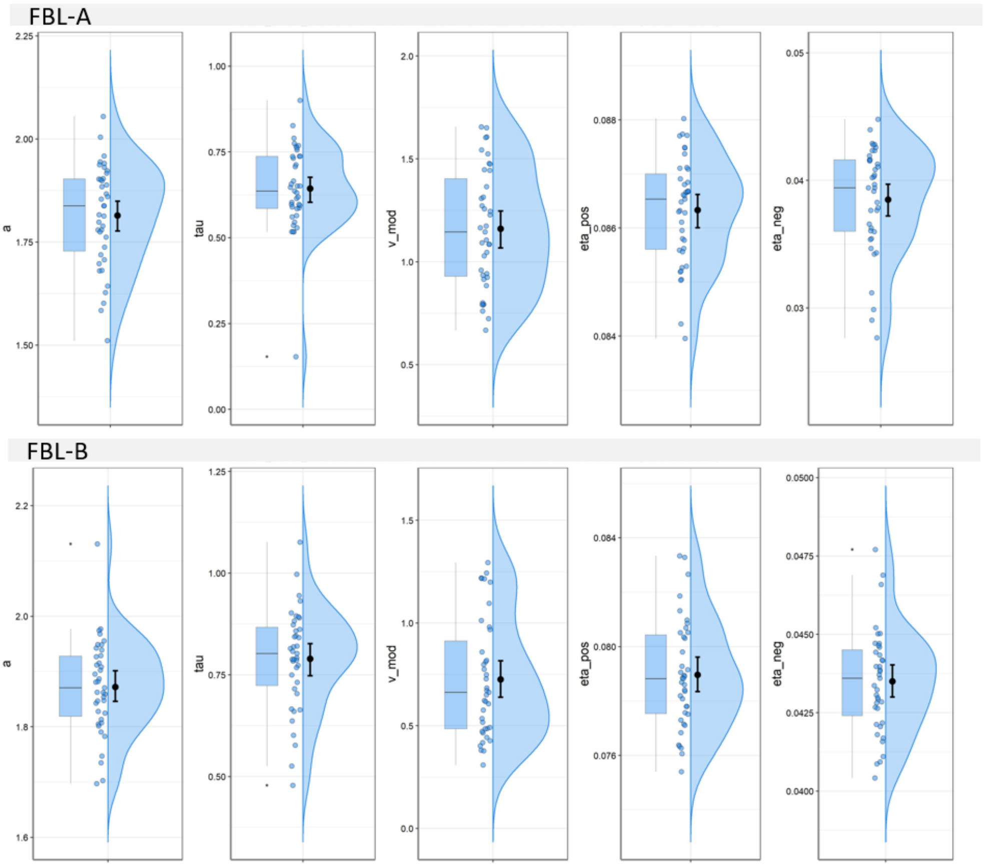

Figure A.5. Distribution of the subject-level model parameters for parts A and B. Error bars represent mean and 95 % CI.

*Correlations between task performance and cognitive skills*

Task performance was assessed by the basic measures (accuracy and reaction times averaged across blocks) and the subject-level parameters derived from the RLDDM (drift rate-$v_{mod}$, decision boundary-$a$, non-decision time- $\tau$ and learning rates $\eta_{+/-}$ ). The main cognitive scores considered were: Non-verbal IQ, SLRT2b, LGTV speed, comprehension and accuracy, RST spelling, digit span, RAN colors and objects (percentile scores were used when available). We found moderate correlations between several cognitive tests (RAN, non-verbal IQ and LGVT text reading comprehension and speed) and task performance, with Pearsons *R* ranging from 0.26 to 0.55. These results are presented in the Supplementary Table A.3.

Supplementary Table A.3. Pearson correlations between task performance and cognitive skills with $p < 0.05$.

| Task performance | Cognitive test | N | *R* | *p* |
|---|---|---|---|---|
| **Basic measures** | | | | |
| FBL A | | | | |
| RT_total | Non-verbal IQ | 32 | -0.55 | .001 |
| RT_total | RAN color (seconds) | 34 | 0.42 | .012 |
| Accuracy_total | Non-verbal IQ | 33 | 0.49 | .004 |
| Accuracy_total | LGVT comprehension PR | 38 | 0.42 | .009 |
| Accuracy_total | LGVT speed PR | 38 | 0.40 | .013 |
| Accuracy_total | RAN color (seconds) | 35 | -0.36 | .032 |
| FBL B | | | | |
| RT_total | RAN object | 35 | 0.40 | .018 |
| Accuracy_total | Non-verbal IQ | 33 | 0.41 | .019 |
| **RLDDM Parameters** | | | | |
| FBL A | | | | |
| $v_{mod}$ | Non-verbal IQ | 33 | 0.49 | .004 |
| $v_{mod}$ | LGVT comprehension PR | 38 | 0.36 | .025 |
| $v_{mod}$ | LGVT speed PR | 38 | 0.36 | .028 |
| $a$ | RAN color (seconds) | 35 | 0.38 | .023 |
| $a$ | Non-verbal IQ | 33 | -0.38 | .029 |
| FBL B | | | | |
| $\eta_+$ | Non-verbal IQ | 33 | 0.39 | .026 |
| $v_{mod}$ | Non-verbal IQ | 33 | 0.37 | .033 |

FBL = feedback learning task; N = sample size after exclusion of outliers (1.5 IQR criterion); PR = percentile score; RT = reaction times; $v_{mod}$ = drift rate modulator; $a$ = decision-boundary; $\tau$ = non-decision time; $\eta_+$ = learning rate for correct responses; RAN = rapid automated naming; LGVT = Lesegeschwindigkeits- und Verständnistest (reading speed and comprehension test)

*Conventional fMRI analysis*

A general linear model (GLM) using individual onsets of stimuli and feedback was convolved with the canonical hemodynamic response function as implemented in SPM12. The onsets of stimuli and feedback presentation were divided into thirds of 16 trials, which resulted into six regressors of interest in the GLM (thirds 1-3, stimuli/feedback onset; as a proxy to describe early learning vs consolidated learning phases in the blocks). The onsets of stimuli and feedback in trials where no responses were given ('too late' responses) were added as an additional regressor of no interest. In addition, six realignment parameters from the data preprocessing were included as nuisance regressors and a binary regressor censored scans with FD > 1 (see MR data acquisition and preprocessing). At the group level, the main effects of interest were the differences between third 3 and third 1 (defining *consolidation* vs *learning* phases) in stimuli and feedback processing. This was tested with one-sample *t* tests using the respective contrast files of each subject. Labels of the resulting brain regions were obtained using the SPM anatomy toolbox [90].

Whole-brain

The whole brain analysis yielded several patterns of activation associated with changes in stimulus and feedback processing along the experimental blocks. These results for each part are presented in Supplementary Table A.4.a. and A.4.b and Supplementary Figure A.6.

*FBL-A. S*timulus processing in the learning phase compared to the consolidation phase (third 1 > third 3) yielded significant activation clusters in right angular and temporal gyrus, as well as the dorsal portions of the caudate and mid frontal areas. In the consolidation phase (third 3 > third 1) stimulus processing resulted in extensive clusters of activations in bilateral occipital regions, including the occipito-temporal fusiform gyrus with a peak in the right fusiform. Additional clusters were found in right insula, right ventral portion of the caudate, and frontal areas. Feedback processing, on the other hand, resulted in stronger activations when comparing the learning vs consolidation phases (third 1 > third 3) in mid frontal areas, cerebellum, right insula, as well as in right mid temporal and angular gyri. In the consolidation phase (third 3 > third 1) clusters of activation were detected in left postcentral, right superior parietal, left mid occipital and temporal parietal cortex, right fusiform and right superior orbitofrontal cortex.

Table A.4a. FBL-A

| contrasts & brain areas | MNI coordinates x | y | z | cluster $p_{FWEcor}$ | voxels | peak Z |
|---|---|---|---|---|---|---|
| *Stimulus onset* | | | | | | |
| third 1 > third 3 | | | | | | |
| Dorsal Caudate R | 22 | 6 | 21 | 0.007 | 83 | 5.14 |
| Frontal Inf Orb R | 43 | 48 | -9 | 0.004 | 93 | 4.97 |
| Angular R | 61 | -60 | 33 | 0.000 | 154 | 4.7 |
| Dorsal Caudate L | -17 | 12 | 21 | 0.019 | 68 | 4.42 |
| Mid Frontal R | 37 | 48 | 30 | 0.015 | 72 | 4.3 |
| Mid Temporal R | 67 | -24 | -9 | 0.000 | 140 | 3.9 |
| third 3 > third 1 | | | | | | |
| Sup Occipital L | -17 | -90 | 3 | 0.000 | 3631 | 5.68 |
| Fusiform R | 31 | 3 | -36 | 0.000 | 933 | 5.44 |
| Insula R | 37 | -6 | 12 | 0.007 | 84 | 4.93 |
| Supp Motor Area R | 7 | -6 | 54 | 0.000 | 339 | 4.56 |
| Postcentral R | 40 | -36 | 72 | 0.000 | 247 | 4.52 |
| Inferior Frontal L | -38 | 15 | 24 | 0.005 | 89 | 4.33 |
| Ventral Caudate R | 13 | 12 | -9 | 0.007 | 85 | 4.23 |
| *Feedback onset* | | | | | | |
| third 1 > third 3 | | | | | | |
| Mid Frontal R | 40 | 45 | 18 | 0.000 | 278 | 4.87 |
| Cerebellum L | -41 | -54 | -45 | 0.009 | 75 | 4.79 |
| Mid Frontal R | 40 | 21 | 42 | 0.010 | 74 | 4.41 |
| Insula R | 37 | 18 | -3 | 0.012 | 72 | 4.31 |
| Angular R | 43 | -51 | 30 | 0.001 | 119 | 4.19 |
| Mid Temporal R | 55 | -21 | -9 | 0.001 | 107 | 3.9 |
| third 3 > third 1 | | | | | | |
| Postcentral_L | -29 | -30 | 75 | 0.000 | 390 | 5.18 |
| Sup Parietal R | 28 | -57 | 75 | 0.007 | 79 | 5.15 |
| Mid Occipital L | -20 | -93 | 3 | 0.000 | 142 | 5.14 |
| Mid Temporal Pole L | -29 | 9 | -33 | 0.003 | 92 | 5.12 |
| Sup Parietal L | -8 | -84 | 60 | 0.013 | 70 | 4.63 |
| Fusiform R | 34 | 0 | -36 | 0.030 | 58 | 4.53 |
| Sup Orbito frontal R | 16 | 48 | -12 | 0.040 | 54 | 3.75 |

Significant activations at whole-brain cluster-level with a cluster defining threshold, $p_{CDT} < .001$; FWEcor = Familywise error-corrected; mid Cingulum = median cingulate gyrus; Sup = superior; Mid = middle; L= left hemisphere; R = right hemisphere

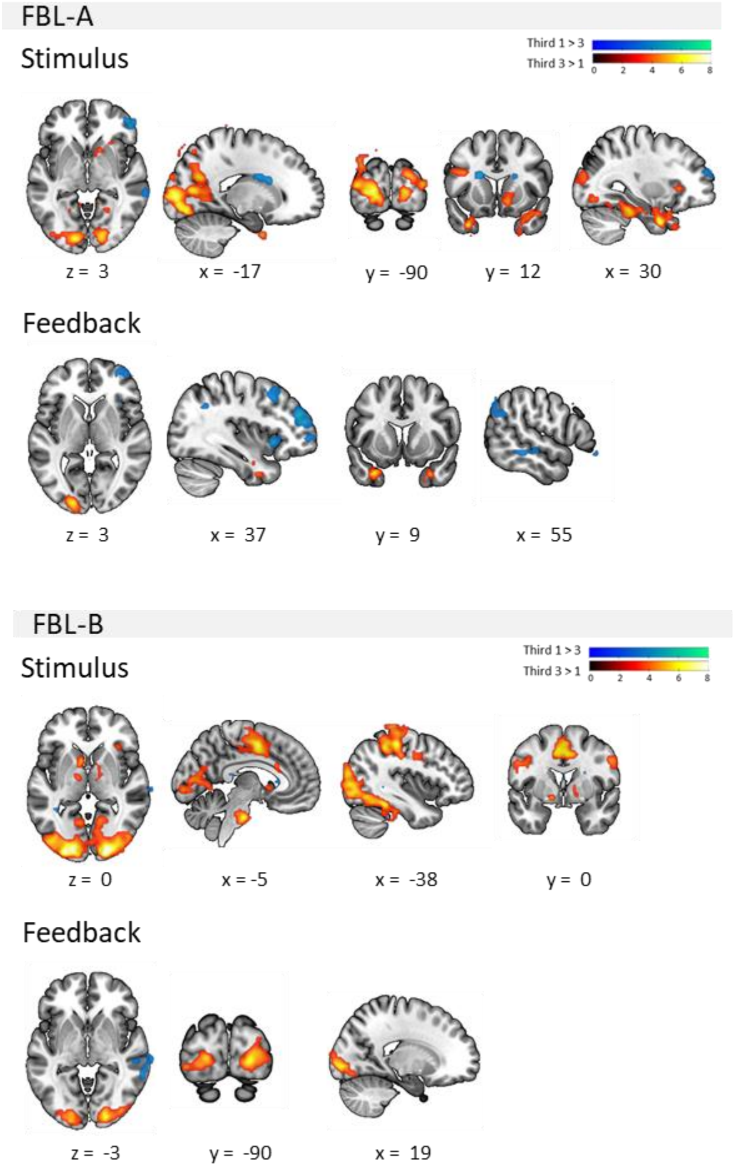

Figure A.6. Suprathreshold activation clusters in FBL-A and FBL-B for stimulus and feedback processing in contrasts comparing the first vs last thirds of trials. Details are provided in Tables 4.a and b. $p_{CDT} < .001$, $p_{FWEc} < .05$. Coordinates in MNI-152 space.

*FBL-B. S*timulus processing in the learning vs consolidation phase (third 1 > third 3) yielded no significant suprathreshold clusters. In the consolidation vs learning contrast (third 3 > third 1) there was a large activation cluster spanning occipital and occipito-temporal regions and a peak of activation at the right Calcarine sulcus. Additionally, activation clusters covered the left and right postcentral gyri, right insula and pallidum and left mid Cingulum, caudate nucleus, superior parietal cortex, and amygdala.

Feedback processing in the learning phase (third 1 > third 3) only resulted in a cluster with a peak in the right mid temporal gyrus. In the consolidation phase (third 3 > third 1) with a region in the right Calcarine and left lingual gyrus were detected.

Table A.4b. FBL-B

| Contrast & brain areas | MNI coordinates x | y | z | cluster $p_{FWEcor}$ | voxels | peak Z |
|---|---|---|---|---|---|---|
| *Stimulus onset* | | | | | | |
| third 3 > third 1 | | | | | | |
| Calcarine R | 22 | -93 | 0 | 0.000 | 3569 | 6.58 |
| Mid Cingulum L | -5 | -3 | 48 | 0.000 | 687 | 5.61 |
| Postcentral L | -38 | -30 | 51 | 0.000 | 1070 | 5.37 |
| Postcentral R | 43 | -21 | 42 | 0.000 | 556 | 5.26 |
| Caudate L | -8 | 12 | -3 | 0.009 | 79 | 4.97 |
| Sup Parietal L | -11 | -84 | 57 | 0.001 | 119 | 4.84 |
| Insula R | 34 | 24 | 6 | 0.015 | 70 | 4.81 |
| Amygdala L | -14 | 0 | -9 | 0.018 | 67 | 4.57 |
| Pallidum R | 13 | 0 | 3 | 0.010 | 77 | 3.99 |
| *Feedback onset* | | | | | | |
| third 1 > third 3 | | | | | | |
| Mid Temporal R | 61 | -27 | -12 | 0.001 | 127 | 3.95 |
| third 3 > third 1 | | | | | | |
| Calcarine R | 19 | -90 | -3 | 0.000 | 340 | 5.28 |
| Lingual L | -17 | -90 | 0 | 0.000 | 279 | 4.76 |

Significant activations at whole-brain cluster-level with a cluster defining threshold, $p_{CDT} < .001$; FWEcor = Familywise error-corrected; mid Cingulum = median cingulate gyrus; Sup = superior; Mid = middle; L= left hemisphere; R = right hemisphere

### ROI

The contrast third 3 > 1 for stimulus and feedback onsets were examined in a set of ROIs defined by meta-analysis (see Statistical analysis) to further specify the key regions showing changes in activation after learning in each task. The principal eigenvariate values of the con-images were extracted for each ROI and tested against zero using t-tests. The results are shown in Supplementary Table A.5 and Supplementary Figure A.7.a and b. The results suggest increased in activation in third 3 compared to third 1 for processing stimuli in several reading network regions: in part A this finding was confined to left fusiform and inferior frontal gyrus, but in the part B there was also involvement of right fusiform and left superior temporal gyrus. Regarding the areas involved in learning/ performance monitoring, there was increased activation in caudate nuclei in both parts, but in part B there were significant effects in additional regions of insula and mid cingulate. For feedback processing, on the other hand, there was decreased activation in the last third compared to the first third of the trials in right caudate, bilateral insula and mid cingulum in FBL-A, but no significant effects were found in FBL-B.

Supplementary Table A.5. ROI t-tests in contrast third 3 > 1

| keyword/ brain region | FBL-A | | | | FBL-B | | | |
|---|---|---|---|---|---|---|---|---|
| | Stimuli | | Feedback | | Stimuli | | Feedback | |
| | *t* | *p* | *t* | *p* | *t* | *p* | *t* | *p* |
| L Fusiform | **3.95** | **< .001** | 0.24 | 0.81 | **4.03** | **< .001** | -0.13 | 0.898 |
| R Fusiform | 1.52 | 0.136 | 0.51 | 0.613 | **3.51** | **0.001** | 0.64 | 0.526 |
| L IFG | **3.76** | **0.001** | 0.24 | 0.811 | **2.89** | **0.006** | 0.58 | 0.567 |
| R IFG | 1.65 | 0.107 | -1.82 | 0.076 | 1.84 | 0.074 | -1.08 | 0.288 |
| L STG | 0.55 | 0.583 | 0.29 | 0.776 | **2.13** | **0.039** | 1.45 | 0.155 |
| R STG | 0.08 | 0.938 | 0.39 | 0.702 | 0.84 | 0.407 | -0.37 | 0.713 |
| L Putamen | **3.72** | **0.001** | 2.03 | 0.049 | **3.4** | **0.002** | -0.05 | 0.963 |
| R Putamen | 0.39 | 0.701 | -0.77 | 0.449 | 1.08 | 0.289 | -1.43 | 0.161 |
| L Hippocampus | -1.39 | 0.173 | 0.74 | 0.464 | -0.35 | 0.732 | 0.65 | 0.518 |
| R Hippocampus | 0.17 | 0.868 | -0.16 | 0.877 | -0.38 | 0.708 | 0.74 | 0.467 |
| L Caudate | **3.34** | **0.002** | 1.71 | 0.095 | **3.42** | **0.002** | 0.03 | 0.977 |
| R Caudate | **3.93** | **< .001** | **2.02** | **0.05** | **2.49** | **0.017** | 0.01 | 0.991 |
| L Insula | 1.74 | 0.090 | **-3.71** | **0.001** | **2.32** | **0.026** | -0.28 | 0.782 |
| R Insula | -0.09 | 0.927 | **-3.5** | **0.001** | **2.77** | **0.009** | -0.84 | 0.404 |
| L mid Cingulum | 0.25 | 0.802 | **-2.63** | **0.012** | **3.03** | **0.004** | 0.17 | 0.865 |
| R mid Cingulum | -1.19 | 0.240 | **-3.08** | **0.004** | **2.91** | **0.006** | -0.2 | 0.843 |

mid cingulum = median cingulate gyrus; left hemisphere; R = right hemisphere; STG= superior temporal gyrus; IFG = inferior frontal gyrus

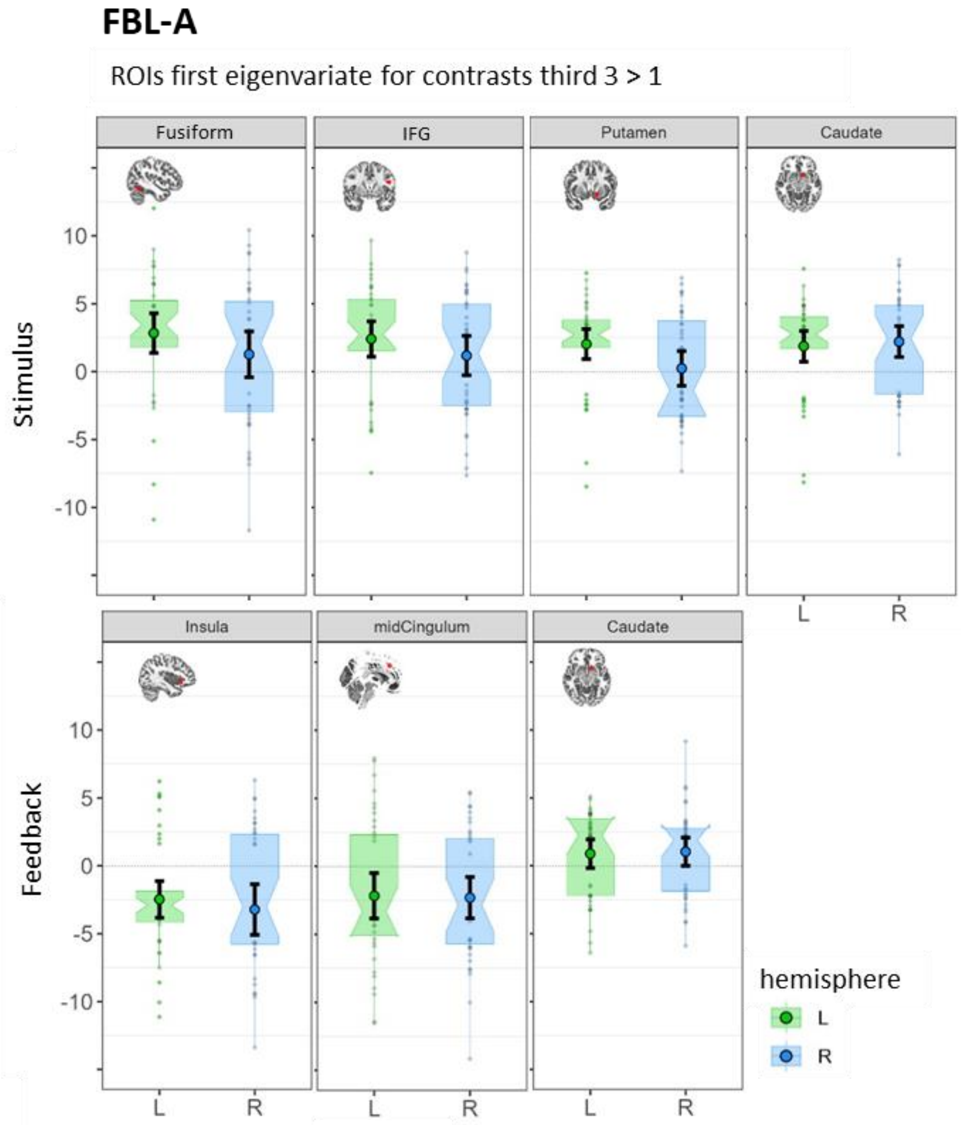

Figure A.7.a. First eigenvariates for the region of interest (ROI) showing significant activations in the contrast third 3>1 for processing stimuli and feedback in FBL-A. Left hemisphere (L) regions are presented in green and their right (R) homologues in blue. Boxplots are notched around the median and error bars represent 95 % CI. The embedded brain images show the corresponding ROI mask.

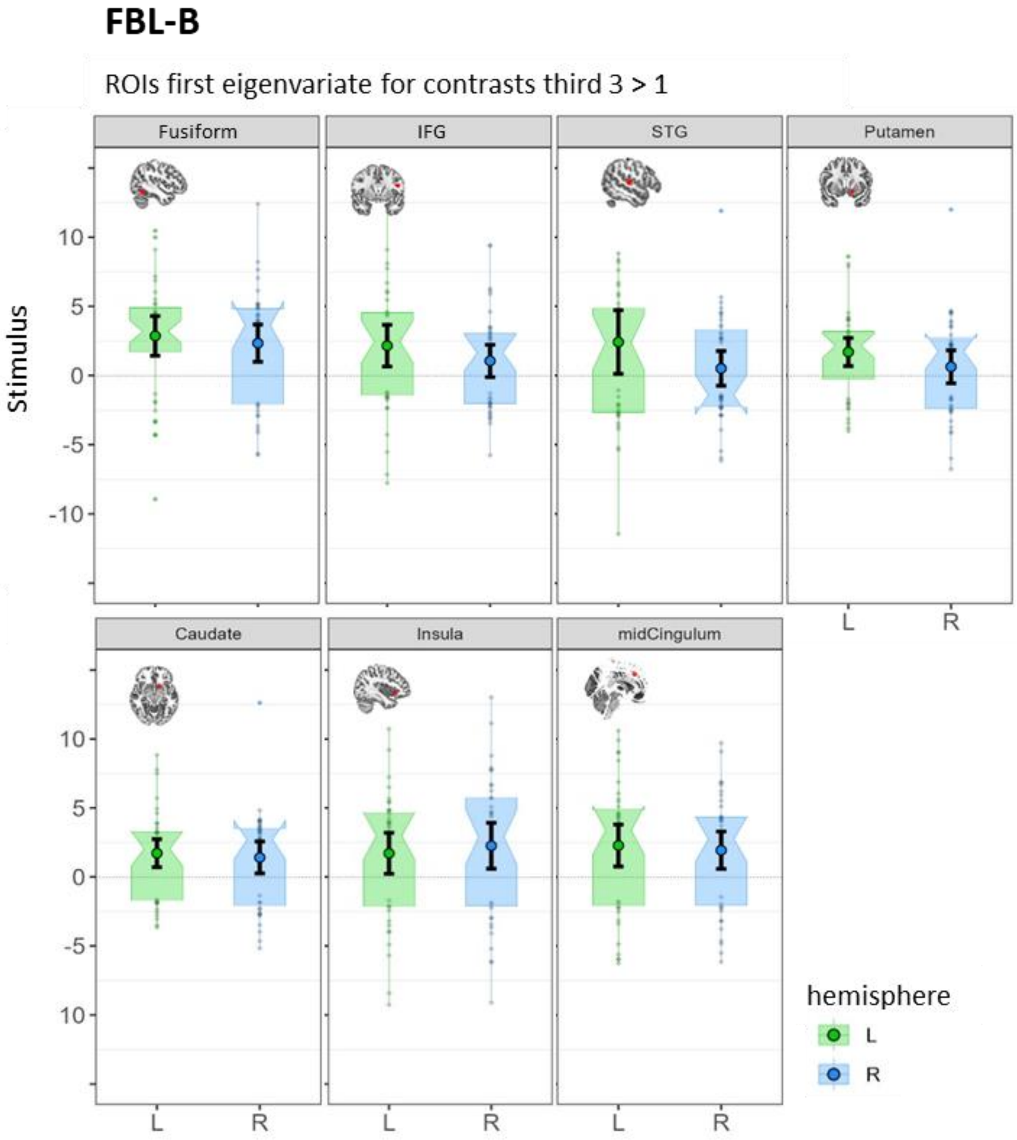

Figure A.7.b. First eigenvariates for the region of interest (ROI) showing significant activations in the contrast third 3>1 for processing stimuli in FBL-B. Left hemisphere (L) regions are presented in green and their right (R) homologues in blue. Boxplots are notched around the median and error bars represent 95 % CI. The embedded brain images show the corresponding ROI mask.
